# A bespoke analytical workflow for the confident identification of sulfopeptides and their discrimination from phosphopeptides

**DOI:** 10.1101/2023.07.15.549150

**Authors:** Leonard A. Daly, Dominic P. Byrne, Simon Perkins, Philip J Brownridge, Euan McDonnell, Andrew R. Jones, Patrick A. Eyers, Claire E. Eyers

## Abstract

Protein tyrosine sulfation (sY) is a post-translational modification (PTM) catalysed by Golgi-resident Tyrosyl Protein SulfoTransferases (TPSTs). Information on protein tyrosine sulfation is currently limited to ∼50 human proteins with only a handful having verified sites of sulfation. The contribution of this chemical moiety for the regulation of biological processes, both inside and outside the cell, remains poorly defined, in large part due to analytical limitations. Mass spectrometry-based proteomics is the method of choice for PTM analysis, but has yet to be applied for the systematic investigation and large-scale analysis of biomolecular sulfation (constituting the ‘sulfome’), primarily due to issues associated with discrimination of sY-from phosphotyrosine (pY)-containing peptides. In this study, we developed a mass spectrometry (MS)-based workflow centred on the characterization of sY-peptides, incorporating optimised Zr^4+^-IMAC and TiO_2_ enrichment strategies. Extensive characterization of a panel of sY- and pY-peptides using an array of MS fragmentation regimes (CID, HCD, EThcC, ETciD, UVPD) highlights differences in the ability to generate site-determining product ions, which can be exploited to differentiate sulfated peptides from nominally isobaric phosphopeptides based on precursor ion neutral loss at low collision energy. Application of our analytical workflow to a HEK-293 cell extracellular secretome facilitated identification of 21 new sulfotyrosine-containing proteins, several of which we validate enzymatically using *in vitro* sulfation assays. This study demonstrates the applicability of our strategy for confident, high-throughput, ‘sulfomics’ studies, and reveals new sY interplay between enzymes relevant to both protein and glycan sulfation.

---

Protein phosphorylation involves the reversible, covalent addition of a phosphate group to the side chain of amino acids, predominantly Ser, Thr and Tyr, alongside 9 non-alcoholic amino acid substrates.^2-4^ As a highly abundant post-translational modification (PTM), phosphorylation has been extensively investigated from both an analytical and functional perspective. It plays critical roles in almost all physiological processes, with an estimated ∼90% of human proteins able to be phosphorylated *in vivo* (the phosphoproteome).^5^ Phosphorylation is catalysed by protein kinases, and >500 enzymes encoded within the human genome have the ability to catalyse the transfer of the γ-phosphate group from adenosine triphosphate (ATP) to amino acid residues on substrates. Approximately 100 of these kinases are tyrosine specific,^6^ while others can phosphorylate Ser/Thr and Tyr, or Ser/Thr alone. In contrast, protein sulfation, *i.e*. the covalent addition of sulfate which is transferred from the 3’-phosphoadenosine-5’-phosphosulfate (PAPS) co-factor, is poorly characterized analytically and is predicted to be considerably less abundant than phosphorylation. Furthermore, unlike phosphorylation, sulfation is believed to be relatively specific for tyrosine residues, being catalyzed in humans by two separate gene products that encode Tyrosyl-Protein-Sulfo-Transferases (TPSTs), termed TPST1 and 2. Some 51 human proteins are annotated in UniProt as containing ‘sulfotyrosine’ (sY), but only 33 of these have been experimentally validated, compared with over 11,000 human “phosphoprotein” entries (accessed May 2023).^7^ Experimental and computational estimates based on site conservation and biological environment suggest that ∼1-7% of all Tyr residues have the potential to be sulfated (in flies and mice),^8, 9^ which potentially makes sY a much more prevalent tyrosine-based PTM than often assumed.

sY was first identified over 50 years ago in fibrinogen/gastrin, and appears to be governed (minimally) by acidic consensus motifs in protein targets, which were first characterized biochemically in the early 1980’s.^10-13^ Sulfation results in biologically-relevant changes in protein:protein affinity, modulating host-pathogen interactions, chemotaxis, FGF7 signalling, proteolytic peptide processing and HIV entry *via* the viral chemokine co-receptor CCR5.^14-22^ Mice lacking TPST1 have defects in body mass and retinal function, whereas TPST2-deficient mice exhibit hyperthyroidism and infertility.^23^ In contrast, double TPST knockout mice have high perinatal mortality rates due to pulmonary asphyxia, and lack detectable sY, confirming that TPST1/2 are together rate-limiting for sY deposition *in vivo*.^23^

While TPST-dependent protein sulfation occurs in the Golgi and is believed to be an irreversible modification, sulfation has been most extensively identified on a variety of extracellular/secreted proteins. Interestingly, a recent study reported cytosolic sY of the tumour suppressor p53,^24^ suggesting that the tyrosine ‘sulfome’ may be more extensive and diverse than previously predicted, revealing potential gaps in our current knowledge that are attributable to sub-optimal analytical capabilities for accurate high-throughput investigation of sY.

Phosphotyrosine (pY) and sY are chemically ‘similar’ with near isobaric mass (sY being 9.6 mDa lighter than pY; pY:79.966331 Da and sY: 79.956815 Da), and are of comparable size and shape, containing at least one negative charge at physiological pH, formally pY = -2, sY = -1, Figure 1A. Despite these similarities, the best-established phosphopeptide enrichment protocols and gas-phase fragmentation strategies are highly unsuitable for sY-peptide isolation and analysis, stymieing isolation and MS-based investigation.^25-29^ Currently, sY characterization is reliant on, and highly restricted by, low-throughput TPST screens *e.g*. on immunoprecipitated putative substrates and/or immunoblotting with monoclonal anti-sY antibodies, ^35^S-radio-labelling or fluorescent-based enzyme assays.^8, 17, 20, 30-38^ While such techniques are useful for confirming prevalent sY sites in simple mixtures, these approaches are restricted to known or predicted (acidic motif) sites of sulfation, which limits their application for discovery purposes. Hence, there is a clear and pressing need for the development of bespoke analytical pipelines that are compatible with high-throughput (HTP) technologies to identify & interrogate sulfomes.

**Figure 1:**
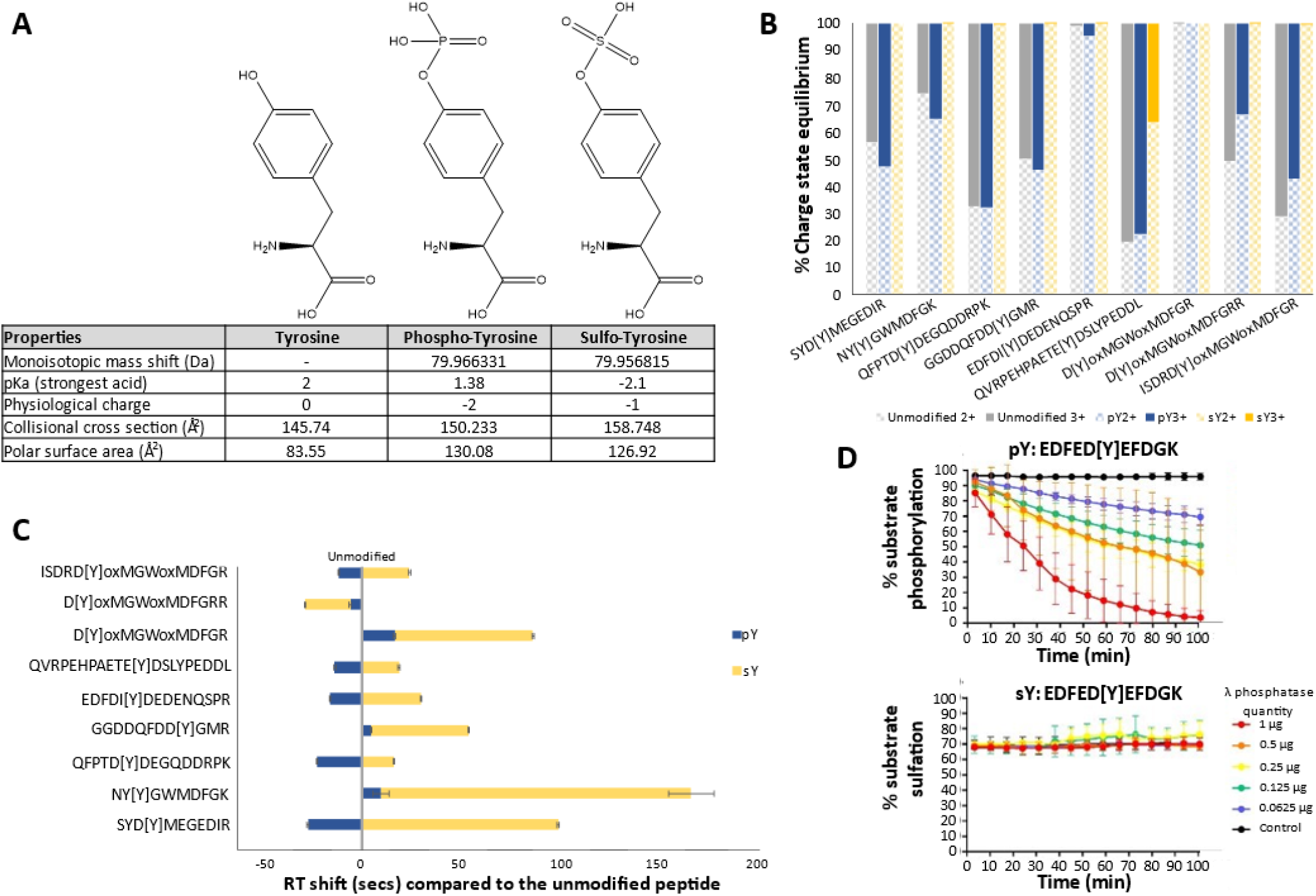
Physiochemical properties of phospho-tyrosine versus sulfo-tyrosine. A) Chemical structure and properties of Tyrosine, Phospho-Tyrosine and Sulfo-Tyrosine. A table of the monoisotopic mass shift (Da) of the phosphate/sulfate moiety in comparison to Tyrosine, the pKa value of the strongest acidic group, net charge influence at physiological pH (∼8), collisional cross section (å) and the polar surface area charge is distributed across (å^2^) are presented. Data were compiled from the 2022 Human Metabolome Database. B) Charge state distribution of our standard panel of peptides in an unmodified, phosphorylated or sulfated state. Modification site denoted by [Y], ox = oxidized residue. N = 10 replicates. C) Scaled retention time shifts of pY- and sY-peptides. Normalised retention time shifts (s) are plotted with reference to the retention time of the unmodified peptide counterpart. Modification site denoted by [Y], ox = oxidized residue. N = 10 replicates. D) Real-time λ phosphatase assay to study peptide dephosphorylation or desulfation. Identical fluorescent peptides containing either pY or sY were incubated with stated quantities of λ phosphatase and the level of modification reversal (dephosphorylation or desulfation) was determined over time (min) using a ratiometric real-time assay.

Recent advancements in mass spectrometry (MS)-based phosphoproteomics pipelines have facilitated interrogation of phosphorylation-dependent signalling networks to an unprecedented level of detail, consolidating it as an essential technique to explore phosphorylation-mediated signalling. An essential part of these analytical pipelines is the selective enrichment of phosphopeptides, based on inherent properties of the appended phosphate moiety, notably a formal net negative charge. Exploitation of standard enrichment strategies, which include titanium dioxide (TiO_2_), anion exchange and Immobilised Metal-ion Affinity Chromatography (IMAC) utilising various metal ions including: Ga^3+^, Ti^4+^ and Fe^3+^, can aid analysis of select sY-peptide standards (typically 2-3 non-tryptic, synthetically sulfated peptides). However, these enrichment processes are reported to be highly inefficient.^25-27^ Compounding this, the relatively low ionization efficiency of sulfopeptides and the high lability of the sulfoester bond during MS analysis compromises sensitivity: neutral loss occurs during collision-induced dissociation (and sometimes during electrospray ionization-MS in the absence of induced fragmentation) generating product ions equivalent to non-modified fragments, hampering sulfosite localization and discrimination from pY-containing peptides.^28, 29, 39^

Electron transfer dissociation (ETD) and related hybrid fragmentation strategies such as EThCD and ETciD (which use supplemental collisional dissociation), have proven useful for characterizing peptides that contain labile PTMs, notably phosphorylation.^40-42^ However, sulfopeptides still remain highly prone to sulfate ‘neutral loss’.^28^ Direct implementation of electron transfer-mediated approaches in a high-throughput (positive ion mode) proteomics study thus do not appear suitable for sensitive sulfation analysis against the relatively small selection of sY peptides so far investigated.

Halim *et al*.^43^ report the application of UVPD at 213 nm (as well as infrared multiphoton dissociation (IRMPD) and high-low photodissociation (HiLoPD)) for sulfopeptide characterisation. In line with the observations of others,^39, 43, 44^ they demonstrate reduced stability of a singly protonated sulfopeptide compared with the doubly charged species. While their data indicated that UVPD at 193 nm (and to a lesser degree, 213 nm) provided almost complete sequence coverage for higher charge state sulfopeptide cations, they reported extensive sulfonate loss.

Negative ion mode MS offers potential advantages for sY-peptide characterization, given the acidic nature of both the sY residue and the side chains surrounding the site of modification reported in most substrates, with reports of greater ionization efficiency and less sulfonate neutral loss, akin to that observed for phosphopeptides.^27, 39, 44^ In particular, negative ETD (NETD) and negative ion mode electron capture dissociation (niECD) show promise for the analysis and localization of peptide sulfation sites, with niECD generating spectra with complete sulfonate retention on the 5 non-tryptic sY-peptides investigated.^44^ Negative ion UVPD also shows promise for localization of sY on peptides, having been used to analyze 5 non-tryptic peptides (4 in common with ^44^). However, optimal characterisation utilises a custom 193 nm laser that is not currently available on commercial instrumentation.^39^

While there are reports of large-scale negative ion mode proteomics studies,^45, 46^ negative ion mode is not routinely applied for HTP studies, in part due to a lack of tested/trained search engines and data analysis tools, as well as reduced commercially available options for fragmentation. Computational tools that are developed for automated interpretation of negative ion mode tandem mass spectra will need to take account of the different and more complex fragmentation pathways that arise *cf* positive ion mode during collision-induced dissociation, including multiple, and extensive, side chain neutral losses.^47-52^

In this paper, we present a proteomics pipeline that can be readily implemented using current commercial instrumentation and search algorithms that is optimised for the HTP identification of sY-containing peptides from both chemically-defined and cellular mixtures, in a manner that permits their discrimination from pY (and other) phosphate-containing peptides. As well as developing a sY-peptide enrichment strategy that employs acetic acid-based solutions with TiO_2_ or IMAC-Zr^4+^, we substantially advance sY-peptide fragmentation studies. Indepth analysis of the largest panel of tryptic sulfopeptides (and phosphopeptide analogues) assembled to date using collision-based, electron-mediated and UVPD fragmentation strategies allowed us to optimise conditions for sY *versus* pY discrimination, exploiting the substantive differences in the energy required for sulfonate *versus* phosphate neutral loss. We also present, and validate, the first application of our sY specific LC-MS/MS-based pipeline for characterisation of the sulfome from a complex cellular mixture. The confident identification of sY-proteins includes 21 novel human sulfoproteins, several of which we validate as TPST1/2 substrates using *in vitro* sulfotransferase assays and MS-based tryptic peptide characterization.

## Experimental Section

### Reagents

Powdered chemical reagents were purchased from Merck. Custom peptides (unmodified and phosphorylated) were purchased from ThermoFisher as PEPOTEC light grade 2 purity. High performance liquid chromatography (HPLC) grade solvents for MS were purchased from ThermoFisher Scientific. All Eppendorf tubes used are ultra-high recovery Eppendorf tubes (STARLAB). PAPS (adenosine 3′-phosphate 5′-phosphosulfate, lithium salt hydrate) was purchased from Merck and stored at -80 °C to afford maximal stability.

### Purification of recombinant proteins

Human TPST1 (residues Lys43–Leu360) and TPST2 (residues Gly43–Leu359) (both lacking the transmembrane domains) were purified as recombinant proteins with an N-terminal 6xHis-tag and a 3C protease cleavage site using pOPINF (OPPF-UK) and refolded *in vitro* into a catalytically-active conformation, as previously described.^31^ Lambda (λ) protein phosphatase was purified to homogeneity as previously described.^53^ The tyrosine kinase EphA3, which autophosphorylates on Tyr when expressed in *E. coli*, comprised the kinase domain and the juxtamembrane region with an N-terminal 6xHis-tag, and was expressed in pLysS *E. coli* from pET28a LIC vector, and purified using Ni-NTA agarose and size exclusion chromatography.^54^

### In-vitro sulfation of peptide standards

Peptide sulfation assays were performed as described previously ^31^. Briefly, 2.5 μg of each peptide in individual reactions were incubated at 20 °C for 18 h in the presence of 1 μg TPST1 and TPST2, and 0.5 mM of the co-factor PAPS, the source of the transferred sulfate. In vitro sulfated peptides were combined before undergoing stage-tip based strong cation exchange clean-up.^55^ The eluent was aliquoted into 0.25 μg/peptide fractions and vacuum-dried by centrifugation prior to analysis.

### Protein phosphatase treatment

For cell-based experiments, a final concentration of 2 mM MnCl_2_ and 1 mM dithiothreitol was added to the sample followed by addition of a 100:1 (w/w) ratio of peptide: purified λ phosphatase. Samples were incubated at 37 °C for 2 hr with constant shaking at 600 rpm.

### Enzymatic sTyr and pTyr peptide modification assays

A 5-FAM fluorophore (maximal excitation of 495 nm/maximal emission of 520 nm, detectable by a LED-induced fluorescence), was covalently coupled to the N-terminus of a custom peptide substrate containing a single modifiable Y site embedded in a canonical acidic motif (5-FAM-EDFED[Y]EFDG-CO-NH_2_) which was purchased from Pepceuticals (Leicester, U.K). The peptide was enzymatically Tyr sulfated as described above, or enzymatically Tyr phosphorylated using the same buffer conditions but substituting TPST1 and 5 mM PAPS with the tyrosine kinase EphA3 and 5 mM Mg:ATP. The PerkinElmer Lab-Chip EZ II Reader system, 12-sipper chip and CR8 coating, and assay separation buffer, were all purchased from PerkinElmer and employed as reported previously.^56, 57^ Pressure and voltage settings were adjusted manually to afford optimal separation of modified (sY and pY) vs. unmodified peptides. Individual phosphatase assays were performed (in triplicate) in a 384-well plate in a volume of 80 μl using 2 μM peptide in the presence of the indicated quantity (μg) of λ phosphatase, 50 mM HEPES (Ph7.4), 0.015% (v/v) Brij-35, 5 mM MnCl_2_ and 1 mM DTT. The degree of peptide sulfation/phosphorylation was directly calculated by the EZ Reader software by differentiating modified : unmodified phosphorylated or sulfated peptide peak ratios.

### Titanium dioxide enrichment

Dried peptide standards were resuspended in the appropriate buffers to a concentration of 0.2 μg/μl by water bath sonication for 10 min. TiO_2_ resin (GL-Sciences) was added to a final amount of 5:1 (w/w) TiO_2_ resin : peptide and incubated at 25 °C with 1,500 rpm shaking for 30 min before centrifugation at 2,000 *g* for 1 min at room temperature and supernatant removed. TiO_2_ resin-peptide complexes were washed 3x for 10 min with 1,500 rpm shaking in the same buffer, resuspending in an equal volume after centrifugation as before. TiO_2_ resin was vacuum centrifuged for 15 min, prior to addition of an equal volume 5% ammonium hydroxide (in water) and shaking at 1,500 rpm for 10 min. Samples were centrifuged as before, supernatant collected and dried to completion under vacuum centrifugation.

### Labelling NTA-agarose coated magnetic beads

PureCube NTA MagBeads (Cube BioTech) were labelled according to the manufacturer’s protocol, scaling to 100 μl resin slurry using magnetic support racks. Pre-labelled Ti^4+^ and Zr^4+^ PureCube MagBeads were washed following the labelling protocol. All beads were made to a 1% (w/v) stock and stored at 4 °C prior to use.

### Immobilized metal affinity chromatography enrichment

An adapted version of the TiO_2_ enrichment protocol above was used. Alterations include: using 2.5:1 ratio of IMAC resin: peptide, and washing the required quantity of beads twice in resuspension buffer using a volume five-fold that of the bead slurry, prior to the addition to peptides.

### C18 stage-tip clean-up

sY-peptides for fragmentation studies, standard input samples and spiked into BSA/Casein digests were subjected to in-house C18 stage-tip (Empore Supelco 47 mm C18 extraction discs)) clean-up prior to LC-MS/MS analysis. Three discs were packed per 200 μL tip and centrifuged at 5000 *g* for 5 min to pack the column. All centrifugation steps were at 4000 *g* for 2 min (or until all liquid has passed through the tip) at room temperature. Stage-tips were equilibrated by sequential washing with 200 μL of methanol, elution buffer (80% (v/v) ACN (acetonitrile) + 0.1% (v/v) TFA in water) and wash buffer (0.1% TFA in water) prior to loading peptide samples in 0.1% TFA. Flow through was re-applied before washing in 200 μL of wash buffer and eluting in 200 μL elution buffer. Elution was dried to completion by vacuum centrifugation.

### Liquid chromatography-tandem mass spectrometry (LC-MS/MS) analysis

All dried peptides were resuspended in 3% (v/v) ACN + 0.1% TFA (in water) by water bath sonication for 10 min, followed by centrifugation at 13000 *g* at 4 °C for 10 min, and the cleared supernatant collected. For protein/peptide standards, peptides were separated by reversed-phase HPLC over a 30 min gradient using an Ultimate 3000 nano system (Dionex), as described in^58^. All data acquisition was performed using a Thermo Orbitrap Fusion Lumos Tribrid mass spectrometer (Thermo Scientific) using stated fragmentation conditions for 2+ to 5+ charge states over a *m/z* range of 350-2000. MS1 spectra were acquired in the Orbitrap [120K resolution at 200 *m/z*], normalized automatic gain control (AGC) = 50%, maximum injection time = 50 ms and an intensity threshold for fragmentation = 2.5e^4^. MS2 spectra were acquired in the Orbitrap [30K resolution at 200 *m/z*], AGC target = normal and maximum injection time = dynamic. A dynamic exclusion window of 10 s was applied at a 10 ppm mass tolerance. For cell-based studies, peptides were separated and acquired as before with either of the following adaptions:1) a 60 min gradient and higher-energy C-trap dissociation (HCD) set at 32% normalized collision energy (NCE). 2) For neutral loss triggered methods, HCD set at 10% NCE, MS2 acquired in the Orbitrap [15K resolution at 200 *m/z*] and targeted loss continuation trigger for *m/z* values equivalent to 1-3 sulfation PTMs (79.9568) at charge states 2+ to 5+ (25 ppm mass tolerance). Continuation of trigger was performed for neutral loss ions with at least 10% relative intensity for correct charge state assigned losses. Triggered scans were acquired in the Orbitrap [30K resolution at 200 *m/z*], HCD set to 32% NCE, AGC target = standard and maximum injection time = auto.

### Mass spectrometry data analysis

The panel of synthetic peptide standards and the products of immunoprecipitated heparan-sulfate 6-O-sulfotransferase 1/2/3 (H6ST-1/2/3) *in vitro* Tyr sulfation assays were analyzed using Proteome Discoverer 2.4 (Thermo Scientific) in conjunction with MASCOT ^59^ against either a custom database of all (12) peptides, BSA and Casein (αS1, αS2, β) isoforms only, or UniProt Human Reviewed database [updated May 2023] for the H6ST immunoprecipitation experiments. Data was imported into Skyline for calculating charge state distributions and retention time shifts based off *m/z* values from Proteome Discoverer 2.4 analysis. Secretome data was converted into MZML format using MSConvert^60^ with peak picking “2-“ filter applied. For neutral loss datasets, an additional filtering parameter of “HCD energy 32” was applied. MZML datafiles were searched using PEAKS11 against the UniProt Human reviewed database (updated June 2022). All data was searched using trypsin (K/R, unless followed by P) with 2 miscleaves permitted and constant modifications = carbamidomethylation (C), variable modification = oxidation (M), sulfation (STY), phosphorylation (STY). Neutral loss triggered methods were additionally searched using semi-specific trypsin and the additional variable modification of deamidation (NQ). MS1 mass tolerance = 10 ppm, MS2 mass tolerance = 0.01 Da, instrument type = Orbitrap (Orbi-Orb), fragmentation = HCD, acquisition = DDA. All data was filtered to 1% FDR.

### Informatics and Localization data analysis

For localization data analysis and heatmap preparation, synthetic peptide MS data sets were searched in Comet (with parameters matched to the PEAKS11 search), which enables export of all PSMs, without any machine learning-based rescoring (as in PEAKS11) or score thresholding. Heat maps were produced in R (tidyverse, ggplot2), for every fragmentation mode (for phosphopeptides and sulfopeptides at multiple charge states). The heat map displays, for correctly identified PSMs, the proportion of identified count of fragment ions containing the known modification site, divided by the total possible count of theoretical observable ions containing the known modification site.

### Cell culture, secretome precipitation and sample preparation

Adherent HEK-293 and HEK-293T cells were seeded at a density of ∼1.75×10^5^ cells/cm^2^ in DMEM media supplemented with 10% (v/v) fetal calf serum, 1% (v/v) non-essential amino acids and 1% (v/v) penicillin/streptomycin and maintained at 37°C, 5% CO_2_ until ∼80% confluent (∼2×10^8^ cells). Cells were washed 2x in 10 ml of phosphate buffered saline (PBS) before incubation for 18 hr in serum-free growth medium. The HEK-293 ‘secretome’ was collected from the medium and centrifuged at 200 *g* for 5 min and then 10,000 *g* for 5 min, preserving the cleared supernatant each time. Secreted proteins were captured by addition of 100:1 (v/v) secretome:Strataclean resin (Agilent Technologies) (∼2 ml) and incubated at room temperature with end over end rotation for 2 h. Strataclean resin-protein complexes were centrifuged at 500 *g* for 2 min and supernatant removed. Complexes were washed twice in an equal volume (relative to initial secretome) of PBS, resuspended in 1 ml PBS and transferred to a 1.5 ml centrifuge tube and washed (3x) 1 ml PBS. Proteins were eluted in 1 ml of 5% (w/v) SDS, 500 mM NaCl and 50 mM Tris (pH 8) at 80 °C for 10 min, with brief vortexing every 2 min, followed by centrifugation at 13,000 *g* for 10 min at room temperature. Protein content was determined by BCA assay before samples were reduced and alkylated with dithiothreitol and iodoacetamide ^58^. A 1:1 mixture of magnetic hydrophobic and hydrophilic Seramag speedbeads (MERCK) were added at a 2:1 (w/w) ratio of beads : protein and ACN was added to a final concentration of 80% (v/v), and incubated at 25°C, shaking (1,500 rpm) for 30 min. The supernatant was discarded and beads washed (3x) in 200 μl of 100% ACN. Beads were dried by vacuum centrifugation for 10 min and resuspended by water bath sonication for 2 min in 1 ml 100 mM ammonium acetate pH 8. Trypsin Gold (Promega) was added at a 33:1 (w/w) protein:trypsin ratio and incubated overnight at 37°C with 1,500 rpm shaking. The peptide containing supernatant was transferred into a fresh tube, and the remaining beads were then washed with a 2x volume of 8 M urea, 100 mM ammonium acetate pH 8 for 30 min with 1,500 rpm shaking, before the supernatants were combined. The pooled sample was incubated on ice for 30 min and centrifuged at 13,000 *g* for 10 min at 4 °C and cleared supernatant collected. Samples were λ phosphatase-treated (as described) prior to acidification with a final concentration of 0.5% (v/v) TFA, and incubated at 37 °C for 30 min followed by 4 °C incubation for 30 min. Samples were centrifuged at 13,000 *g* for 10 min at 4 °C, and the cleared supernatants collected. Samples were split 1:33:33:33 respectively, for total protein, agarose coated IMAC-Zr^4+^, BioResyn IMAC-Zr^4+^ HP or TiO_2_ sY enrichment protocols, prior to drying to completion by vacuum centrifugation.

### Functional enrichment analysis

DAVID Bioinformatics Resources [v6.8]^61^ was used to understand cellular compartments (and biological processes) in the secretome enriched protein sample.

### Heparan-sulfate 6-O-sulfotransferase 1/2 immuno-precipitation and *in vitro* sulfation assasy

The cytoplasmic regions of human HS6ST1 (37-410), HS6ST (228-605), and HS6ST3 (28-471) were cloned into pcDNA3 with a 3C-protease cleavable, N-terminal, tandem StrepTag for human cell expression. HEK-293T cells were transfected at ∼40 % confluency using a 3:1 polyethylenimine (branched, average Mw ∼25,000 Da; Merck) to DNA ratio (30:10 μg, for a single 10-cm culture dish). Proteins (from 10x 10-cm culture dish) were pooled and harvested 48 h post transfection in lysis buffer (150 μl per dish) containing 50 mM tris-HCl (pH 7.4), 150 mM NaCl, 0.5% (v/v) Triton X-100, 2 mM CaCl2, 10 mM MgCl2 and 20% (v/v) glycerol and supplemented with protease and phosphatase inhibitors (Roche). Lysates were sonicated briefly on ice and clarified by centrifuged at 20,817*g* for 20 min at 4 °C, and the resulting supernatants were incubated with 15 μl Strep-Tactin sepharose resin (Thermo Fisher Scientific) for 3 h with gentle end over end mixing at 4 °C. Sepharose beads containing bound protein were collected and washed three times in 50 mM tris-HCl (pH 7.4) and 500 mM NaCl and then equilibrated in storage buffer (50 mM Tris-HCl (pH 7.4), 100 mM NaCl, and 5% (v/v) glycerol). HS6STs were then proteolytically eluted from the beads over a 2 h period using 3C protease (0.5 μg) at 4 °C with gentle agitation. Purified HS6STs (10 μl of the 35 μl elution volume) were sulfated with TPST1, TPST2 or both TPST1 and 2 (as described above) for 18 h at 20 °C.

## Results and Discussion

### Sulfotyrosine and phosphotyrosine have distinct biochemical and analytical properties

The isobaric masses of sY and pY moieties are extremely similar, but possess distinct chemical properties (Fig. 1A; ^62^). At physiological pH (∼7.4), sY carries a single net charge of -1, while pY has a net charge of -2. The primary (most acidic) pKa of sY is -2.1 compared with 1.38 for pY (as reported in the Human Metabolome database (^63^; www.hmdb.ca)). Based on molecular dynamics simulations and experimental analysis, sY exhibits a marked reduction in the potential for hydrogen bonding interactions compared with pY.^62^ This difference in electronegativity likely explains the previously reported reduced ionization efficiency of sY-peptides in positive ion mode and the lability of the sulfonate group during proton-driven collision-induced dissociation ^28, 29, 39^.

To investigate how sY or pY affects peptide analysis, we synthesised a panel of 12 Tyr-containing peptides designed based upon known/putative sulfation sites from proteins. We also generated an analogous panel containing pY. The panel of peptides were enzymatically sulfated using TPST1 and 2 ^31^ to generate a panel of tryptic sY-peptides for comparison against synthetically generated phosphopeptides modified on the same residue (Supp. Table 1, Fig. 1A). All peptide variants (either unmodified, pY- or sY-containing) were analysed under standard positive ion mode LC-MS/MS conditions, comparing charge states and retention time (Fig. 1B, C). Peptides identified in all forms (unmodified, pY and sY) across 10 replicates were included in these analyses (a total of 9 peptides). We identified little difference in the relative abundance of the 2+ and 3+ charge states for peptides when comparing the unmodified and pY forms (Fig. 1B). In marked contrast, and consistent with previous studies,^39, 64^ we observed a significant reduction in the formation of 3+ sY-peptide ions, preferentially observing doubly protonated species; only 1 out of the 9 peptides in this set (QVRPEHPAETE[sY]DSLYPEDDL) presented in its [M+3H]^3+^ form (likely due to the presence of internal Arg and His residues) with all others appearing almost exclusively as 2+ species. Singly protonated sY peptide ions, when present, were at substantially lower levels (<1% relative intensity compared to 2+ ions) suggesting that they are highly unstable.^39, 43, 44^ Comparing LC retention times (RT) for sY-peptides *versus* their unmodified counterparts over the rapid 10 min LC gradient, we observe a clear increase (ranging from 16 s to 156 s *i.e*. a RT shift of between 0.8% and 4.3%) for 8 out of the 9 sY-peptides. Conversely, and as previously reported, pY-peptides exhibited faster LC elution times (ranging from 5 s to 26 s) (Fig 1C).^65, 66^ Interestingly, D[Y]oxMGWoxMDFGRR was the only peptide where RT decreased due to sulfation, and indeed eluted earlier than the pY variant (−22 s for sY versus -5 s for pY). This peptide has the highest pI value of peptides in this panel (pI = 5.96, range= 3.62-5.83, Supp. Table 1) and contains 2 Arg residues, suggesting that there is not a direct relationship between pI and modification type, and hydrophobicity.

Given the comparatively high prevalence of phosphorylated proteins relative to known sulfoproteins in the human proteome, we explored whether enzymatic phosphatase pre-treatment could reduce the relative abundance of phosphopeptides and thereby i) minimize possible mis-identification of sY-peptides, particularly given the very small mass shift between the two moieties (∼9.6 mDa), and ii) improve the sensitivity of sulfoproteomics pipelines by minimizing co-enrichment of sulfopeptides and phosphopeptides. Protein phosphatase treatment combined with MS has previously been used to investigate sY-peptide identification, discriminating from pY due to a lack of activity and thus an absence of mass shift post treatment.^67, 68^ We confirmed the specificity for dephosphorylation by monitoring activity towards pY- and sY-forms of the same fluorescently labelled peptide substrate (EDFED[Y]EFDGK), using a phosphorylation assay ^31^ in reverse (Fig. 1D) similar to the procedure we developed for real-time desulfation analysis of glycans.^69^ Irrespective of the quantity of protein phosphatase employed (μg) and under conditions with essentially complete dephosphorylation of the pY-peptide (>95%), no loss of the sY-containing peptide was observed, confirming that protein phosphatase pretreatment is likely an effective means of reducing phosphopeptide content in biological samples, whilst preserving peptide sulfation prior to enrichment and MS analysis.

### Acetic acid-based solutions with TiO_2_ or Zirconium ^4+^ IMAC can efficiently (and semi-preferentially) enrich sY-peptides

MS-based characterization of PTMs in complex mixtures requires enrichment of peptides (or proteins) to improve detection sensitivity and overcome the fact that PTM modified peptides usually represent a small proportion of the total peptide pool.^2, 70-72^ The selectivity of phosphate-enrichment techniques is based on PTM-specific biochemical properties, or bespoke antibodies. Low pH Immobilised Metal ion Affinity Chromatography (IMAC) and TiO_2_ have thus become staples in phosphoproteomics pipelines.^73-75^ Both TiO_2_ and IMAC-based phosphopeptide enrichment strategies exploit the relatively low pKa of the phosphate group (the primary pKa value of phosphate monoesters being ∼2) compared with the side chains of other amino acids (the most comparable being Asp and Glu with pKa values of ∼3.5 and 4.2 respectively ^76^) to promote efficient and (relatively) specific binding. By maintaining a low pH, it is therefore possible to preferentially enrich for phosphopeptides as opposed to Asp/Glu-rich peptides with positively charged immobilized metal ions.

However, application of these approaches for the enrichment of sY-peptides is potentially complicated by two major factors; firstly, it is reported that sY undergoes acid-induced hydrolysis,^77, 78^ which would reduce the recovery of sY using standard low pH enrichment conditions. However, we failed to detect acid-induced sulfate hydrolysis across our peptide panel, even after sample storage for a week at 4 °C in 0.1% (v/v) TFA (data not shown), in-line with other reports.^27^ Second, the single hydroxyl group of sY and the size/orientation of the sulfonyl group (Fig. 1A) might prohibit the efficient capture of sY-peptides by TiO_2_. Phosphopeptide enrichment with TiO_2_ relies on the spatial coordination and bidentate hydrogen bonding of the phosphate group to resin-chelated titanium ions;^79-81^ the reduced hydrogen bonding capacity of sY-peptides may compromise TiO_2_ binding. Indeed, a previous report that used a commercial TiO_2_ kit reported no enrichment of two sY-peptides.^27^ However, the binding and wash buffers used in this kit are proprietary, and buffer composition is known to play a significant role in the efficiency of such enrichment processes.^82, 83^ Moreover, although sTyr antibodies exist, these are unreliable in our hands at immunoprecipitating proteins or peptides containing sTyr.

While Fe^3+^-IMAC ^27^ and Ga^3+^-IMAC ^26^ have been tested for the enrichment of sY-peptides, the relative efficiency has not been carefully evaluated. Other metal counter ions have also proven useful for phosphopeptide enrichment. ^84-86^ However, there has not been a comprehensive evaluation of the utility of different IMAC counter ions for sY-peptide enrichment. Previous attempts at sY peptide enrichment did not consider tryptic peptides, or fully explore the efficiency or selectivity of sY peptide enrichment as a function of binding and wash conditions. Given our desire to advance sTyr analysis in a variety of biological mixtures, we undertook a comprehensive quantitative evaluation of the ability of different immobilised media, including TiO_2_ and IMAC (comparing 10 different metal counter ions), in terms of specificity and efficiency for enriching tryptic sY-peptides (Figs. 2 & 3).

**Figure 2:**
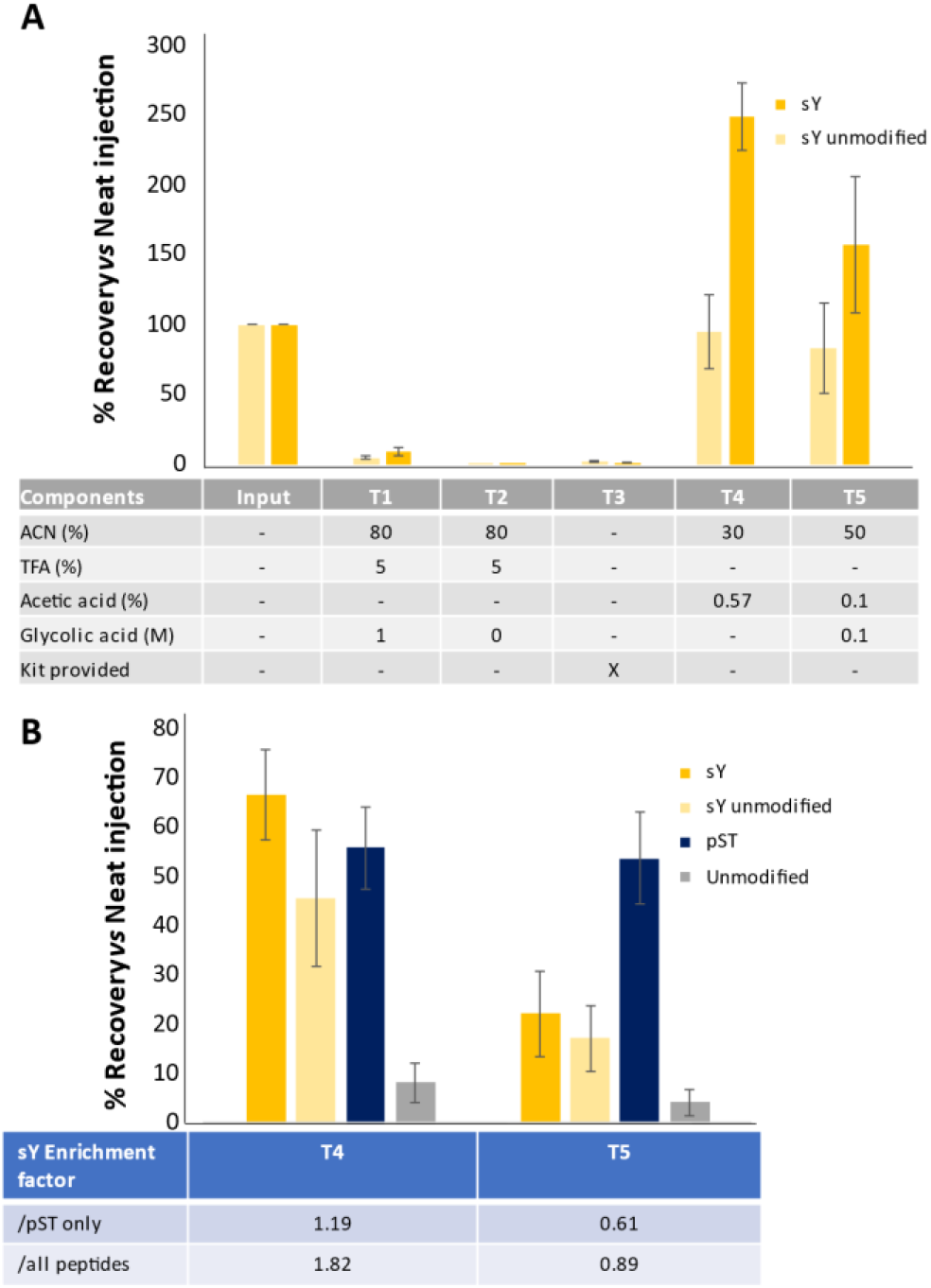
TiO_2_ based enrichment of sY-peptides. A) Solution optimisation for recovery of sY-peptides. The panel of sY- and unmodified peptides were enriched using standard TiO_2_ protocols with different loading and wash solutions as stated and analysed by LC-MS. Recovery was determined by normalising to an equal amount of ‘unenriched’ input material subjected to C18 reverse phase clean-up. Recovery of the 8 sY- and 9 unmodified peptides is an average (mean +/-S.D.) of all equivalent (non-)modified peptides, including methionine oxidised variants which we have previously shown has little effect on relative signal intensity^1^. B) A mixture of trypsin digested BSA, casein-αS1, -αS2 and -β and the sY-peptide panel were subjected to TiO_2_ enrichment using T4 and T5 as specified in (A) and efficiency of enrichment of sY, pS/T and unmodified peptides determined. All samples were subjected to C18 clean-up prior to LC-MS/MS analysis. Recovery (mean +/-S.D.) was determined for 8 sY-, 9 unmodified synthetic peptides, as well as 9 pST peptides from casein, and 19 unmodified peptides from BSA and casein.

Initial evaluations focused on TiO_2_-based enrichment using solutions typically employed for phosphopeptides, and sample loading conditions previously shown to enable binding of three (non-tryptic) sY-peptides ^87^ (Fig. 2). To rule out non-specific enrichment resulting from the increased hydrophobicity of the sY modification, sY peptide recovery rates for each condition were directly compared to the amount loaded following C18 reverse phase chromatography as a ‘clean-up’ step. Under standard TiO_2_ phosphopeptide enrichment conditions (80% ACN, 5% TFA, 1 M Glycolic acid, T1, ^58^), application of a modified loading buffer without glycolic acid (80% ACN, 5% TFA, T2) or using the commercially available Phos-TiO_2_ spin-tip kit (GL Sciences, T3) we failed to observe efficient recovery or enrichment (<5%) of any sY-containing peptides (Fig. 2A). Little to no recovery was observed for non-modified peptides as would be expected. Reducing the acetonitrile content and/or replacing TFA with lower concentrations of acetic acid (30% ACN, 100 mM acetic acid: T4; 50% ACN, 0.1% acetic acid, 0.1 M glycolic acid: T5) improved recovery of all peptides with ∼2.5-fold and 1.5-fold enrichment of sY-peptides for conditions T4 and T5 respectively compared with the C18-desalted control (Fig. 2A).

To evaluate sY-enrichment in a mixture of tryptic synthetic sY peptides alongside phosphopeptides derived from purified BSA and α/β casein (at molar excesses of 170x and 90x respectively), we compared the relative enrichment of sY-versus pST-peptides in these samples. From this peptide mixture we quantified recovery of sY peptides, unmodified peptides (lacking sTyr but still highly acidic), 9 phosphopeptides from casein, and 19 unmodified peptides (10 from BSA, and 9 from casein). Signal intensity was normalised to an equal load of non TiO_2_-enriched material, all samples having been subjected to C18 cleanup (Fig. 2B). sY enrichment factor was then determined by comparing the relative recovery of sY-peptides with respect to either all peptides observed, or phosphopeptides only.

As previously observed for the synthetic peptides (Fig. 2A), sY peptide recovery from this mixture was much lower with the T5 condition than T4, while there was little variation (± <5%) in the relative recovery of either phosphorylated or non-phosphorylated peptides from BSA/casein. Consequently, a greater enrichment factor was observed for T4 than T5, whether normalised against all peptides (1.82 or 0.89 respectively), or just the phosphopeptide cohorts (1.19 or 0.61 respectively). While these results may be explained in part by the reduced acetonitrile content of T4, we hypothesise that the glycolic acid may be acting to reduce non-specific binding of the acidic residues in the sY peptides given the comparative recovery of the unmodified synthetic peptides. Thus, we were able to partially enrich for sY-peptides, and believe that this is primarily due to the high acidic content of TPST1/2-consensus containing peptides.

We next investigated sY-peptide enrichment using IMAC and a panel of 10 metal counter ions (Al^3+^, Co^2+^, Cu^2+^, Fe^2+^, Ga^3+^, Mn^2+^, Ni^2+^, Ti^4+^, Zn^2+^ and Zr^4+^). Under relatively mild conditions (50% ACN, 0.1% TFA), Zr^4+^ exhibited by far the most efficient recovery (∼180 % *cf* C18 enrichment) across all sulfopeptide standards, with Ti^4+^ being the only other counter ion capable of enriching multiple sulfopeptides, albeit rather inefficiently, with a recovery of ∼20% (Fig. 3A). In contrast with previously published data which used Ga^3+^ and Fe^3+^,^26, 27^ we did not observe sulfopeptide capture using Ga^3+^-IMAC and only minimal recovery (∼3%) was seen with Fe^3+^-IMAC.

**Figure 3:**
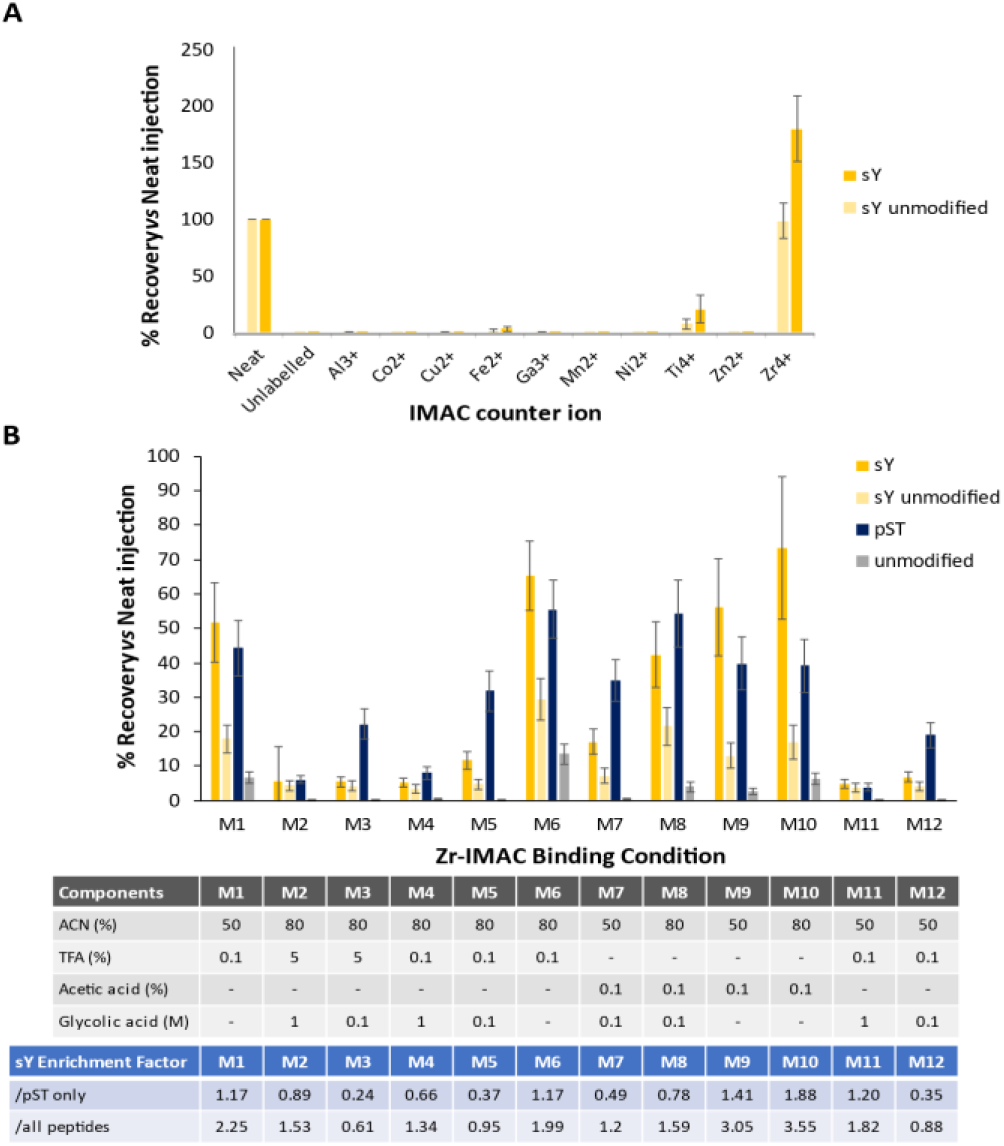
Development of an IMAC based enrichment protocol for sY-peptides. A) Efficiency of recovery of the panel of sY- and unmodified peptides was evaluated for 10 IMAC counterions as indicated. Recovery was determined by normalising to an equal injection of ‘unenriched’ input material subjected to C18 reversed phase clean up. Recovery of the 8 sY- and 9 unmodified peptides is an average (mean +/-S.D.) of all equivalent (non-)modified peptides, including methionine oxidised variants which we have previously shown has little effect on relative signal intensity^1^. B) Optimisation of Zr^4+^ recovery of sY-peptides. A mixture of trypsin digested BSA, casein-αS1/αS2/β and the sY-peptide panel were subjected to enrichment with Zr^4+^ IMAC using different loading solutions as indicated. All samples were subjected to C18 clean-up prior to LC-MS/MS analysis. Recovery (mean +/-S.D.) was determined for 8 sY-, 9 unmodified synthetic peptides, as well as 9 pST peptides from casein, and 19 unmodified peptides from BSA and casein.

Given the comparatively high recovery of sY peptides with Zr^4+^-IMAC, we evaluated the effect of different binding conditions on the recovery and enrichment of sY peptides in a more complex BSA/casein/synthetic peptides mixture, as performed above for TiO_2_. Overall, we evaluated 12 different conditions, altering the concentration of ACN (50%, 80%), TFA (0, 0.1%, 5%), acetic acid (0 or 0.1%) and/or glycolic acid (0, 0.1 M, 1 M) (Fig. 3B). Optimal sulfopeptide recovery with Zr^4+^-IMAC was obtained in the presence of 80% ACN, 0.1 % acetic acid (M10) (∼74% ± 20%, outperforming optimal TiO_2_ conditions). Notably, the relative recovery of phosphopeptides (∼39%) and sY-unmodified peptide (∼17%) was also much reduced compared with optimal TiO_2_-enrichment conditions (Fig. 2B), resulting in enrichment factors of either 3.55 or 1.88 when considering either all peptides, or specificity with regard to phosphopeptide enrichment.

Increasing the acetonitrile concentration (from 50% to 80%) consistently increased the efficiency of sY-peptide recovery (*cf* M7/M8, M9/M10, M4/M11, M5/M12). However, the type and concentration of acid had a greater effect on the efficiency of enrichment, with acetic acid being preferential for sY and TFA being preferential for pY-peptides (M7/M8, M9/M10 versus M4/M11, M5/M12). We also observed a reduction in sY enrichment factors with a decrease in glycolic acid concentration (from 1 M to 0.1 M), primarily due to an increase in the recovery of unwanted phosphopeptides (compare M2/M3, M4/M5, M11/M12). That being said, the overall recovery of sY peptides was substantially reduced in the presence of any concentration of glycolic acid, negating the potentially positive effect on coenrichment of phosphopeptides (M4/M6, M1/M11, M7/M9, M8/M10) (Fig. 3B).

### Challenges associated with accurate site localization of sulfotyrosine within tryptic peptides

While there have been a number of studies exploring different strategies for sulfopeptide characterisation and site localisation (positive *versus* negative ion mode; fragmentation),^27, 28, 39, 43, 44, 64, 67, 68, 88, 89^ the overall utility of these findings in terms of applicability for HTP proteomics studies remains unknown; they either rely on instrumentation not commercially available, or focus on a very small number of non-tryptic peptides and are thus not representative of typical proteomics samples.

To better understand the ability to confidently localize sY sites in tryptic peptides (*i.e*. the generation of site-determining product ions), we characterised product ions generated from our synthetic panel of 12 sY tryptic peptides derived from known protein modifications using all potential fragmentation regimes available on a standard Fusion Lumos Tribrid instrument (Thermo Fisher), namely HCD, CID, ETD, EThcD, ETciD and UVPD. We also evaluated 12 analogous pY peptides, allowing us to compare site localisation confidence for both covalent modifications side-by-side. Considering the multiple settings available for each fragmentation regime, we thus investigated a total of 43 fragmentation conditions (Fig. 4). sY and pY peptide libraries were analysed by LC-MS/MS as separate pools, and searched with COMET to aid analysis. For each fragmentation condition, peptide and charge state, the tandem mass spectrum with the highest COMET score was subjected to further analysis, quantifying the number of observed product ions retaining the covalent modification as a function of the number of theoretically possible product ions (Fig. 4). Unambiguous site localisation ideally requires the generation of product ions that retain the covalent PTM. This is particularly important for sulfation, given the propensity for neutral loss of Δ80 Da resulting in ions that cannot be distinguished from those that would otherwise be unmodified. Data for ion types correlating with fragmentation around a particular residue (*e.g*. a_4_/b_4_/y_(n-4)_, y_8_/y_8_ ^2+/^z_8_) was compiled and treated as a single entity for the purpose of site localisation.

**Figure 4:**
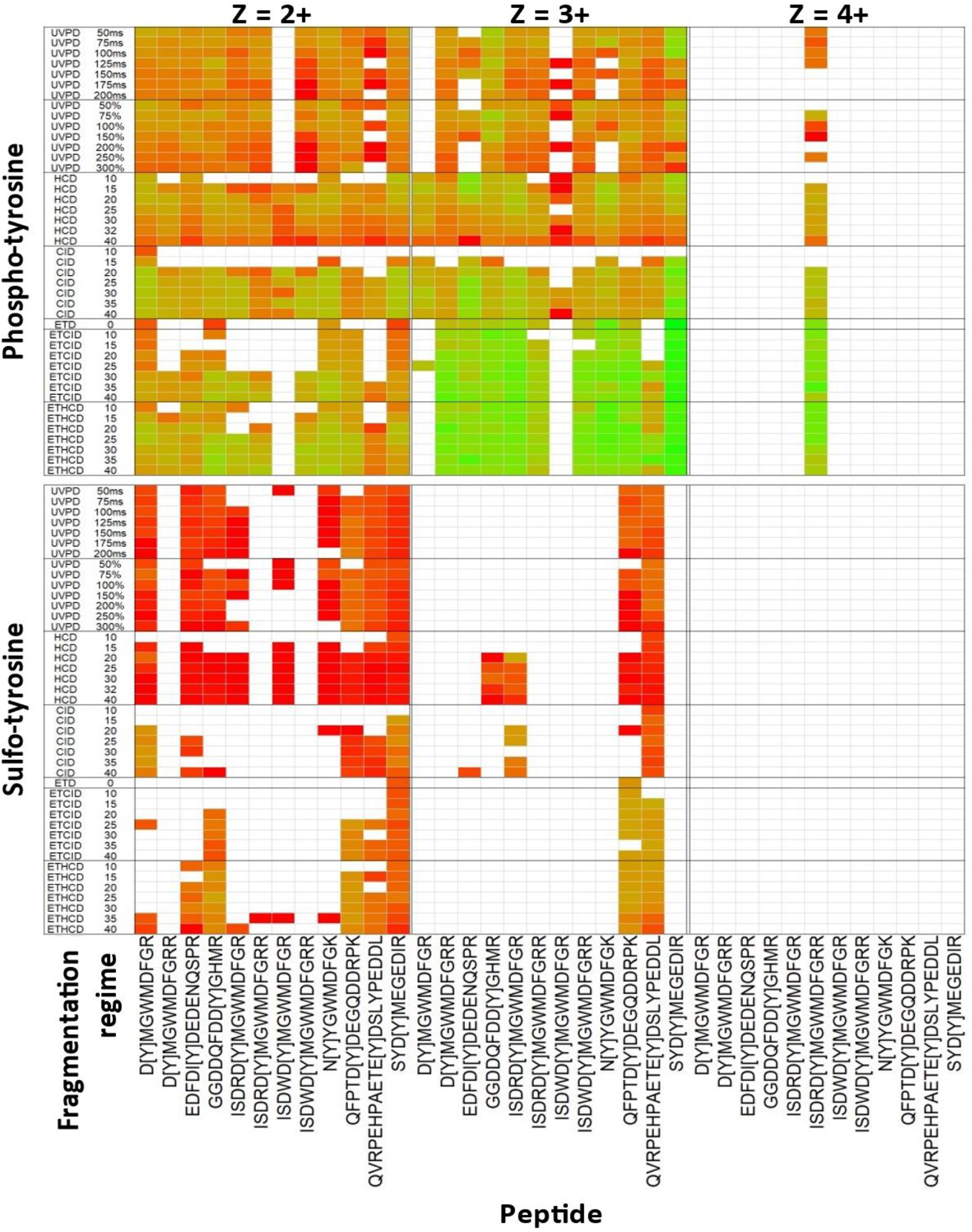
Comparison of site determining ions for sY and pY peptide panel using different fragmentation regimes. The panel of 12 sY- and pY-peptides pools were subjected to LC-MS/MS on the Fusion Lumos Tribrid mass spectrometer (Thermo Fisher) using fragmentation conditions as detailed. Ranging normalised collision energies (NCE) were applied for CID, HCD, ETciD and EThcD as denoted by ‘_X’ (where X = NCE %). ETD component was charge state calibrated. UVPD activation time was either calibrated to molecular weight (%) or manually-set (ms). Data were searched with COMET and the highest scoring PSM selected for further investigation. Heatmap shows log2 ratio of the number of observed MS2 product ions containing the known modification mass shift (Δ80 Da) normalised to the total number of potential PTM-containing product ions. Ions correlating to fragmentation at the same position in the peptide (e.g. a_4_/b_4)_, y_8_/y ^2+^/z_8_) were collapsed into a single entry. Modification site is depicted by ‘[Y]’, and charge states are visualised separately. Green = all potential mass shift containing product ions detected; red = no potential mass shift containing product ions detected, white = no MS/MS spectra were confidently identified.

As shown in Fig. 4, there is stark disparity in the ability to localise the sites of modification on pY-versus sY-peptides, under all the conditions evaluated. Irrespective of the fragmentation strategy employed, and the energetics/time used, very few sulfonate-retaining product ions were observed for sY containing-peptides. ET(hc/ci)D yielded a higher proportion of sY- (and pY-) site determining ions than HCD for those peptides/conditions where product ions were observed, making electron-mediated fragmentation (EThcD at 25% NCE) optimal for localisation of the sY sites in this peptide panel. However, even for these peptides, the relative proportion of site-determining ions was substantially lower than for the pY-peptide equivalents and it is likely that the reduction in sY-peptide ion charge state (Fig. 1B, Fig. 4) prohibited efficient fragmentation (and thus identification) by EThcD/ETciD.

In contrast with HCD, very few sulfopeptides were identified following CID. Where they were identified, there was almost complete neutral loss of sulfonate, with the exceptions of D[Y]MGWMDFGR and SYD[Y]MEGEDIR. While HCD permitted sulfopeptide identification (for 9/12 peptides), the complete loss of 80 Da, even at low NCE (15%), meant that no site determining ions were observed.

Our phosphopeptide data contrasts significantly with that observed for the equivalent sulfopeptides, with varying degrees of phosphate retention seen for all peptides and charge states across all fragmentation strategies employed (Fig. 4). For doubly protonated pY-peptide ions, overall phosphate retention was highest with CID_35, or EThcD_25, while the equivalent 3+ ions benefitted substantially from ET(hc/ci)D, a finding well supported for phosphopeptides in previous studies.^42, 58^ Both fragmentation regimes generally outperformed HCD in terms of the relative proportion of phosphopeptide-retaining product ions across all charge states. Interestingly, our observations with UVPD in terms of both automated peptide identification and generation of site localizing ions broadly mirrored the results with HCD for both sY- and pY-peptides. However, UVPD was highly inefficient for peptide fragmentation, requiring long irradiation times (>100 ms) and generating product ions of low intensity, irrespective of peptide sequence, charge or modifications status. sY-peptides were generally well identified with UVPD (8 out of 12 sY-peptides). However, sulfonate loss was extensive and it was not possible to localise the precise sites of sulfation on our sY-peptide panel.

### sY- and pY-peptides can be discriminated based on precursor neutral loss with HCD at low energy (10% NCE)

A number of strategies have been introduced to help distinguish sY from pY in peptides and proteins. These include a subtractive approach based on phosphatase treatment and acetylation of free (unmodified) tyrosine residues.^67^ However, subtractive analytic approaches fundamentally rely on both complete phosphate removal and subsequent chemical modification of unmodified tyrosine residues, where inefficiencies in either step will result in mis-identification. The presence of additional labile tyrosine PTMs (such as nitration) also increases the potential of sY mis-identifications. Adduction of sY with trace metal ions (from LC solvents and ESI emitters) has also been reported to improve sY peptide identification and site localisation.^64^ As well as increasing the average charge state (thus increasing ETD/EThcD efficiency), metal adducted sY-peptides have been reported to be more likely to retain the sulfate moiety (permitting localization by ETD mediated fragmentation regimes). However, while the relative proportion of Na^+^/K^+^ adducts of the 2 sY-peptides reported in a previous study were relatively high, we failed to observe any sY-peptide metal-ion adduction following manual interrogation and open PTM searching of our data. Unfortunately, addition of Na^+^/K^+^ salts into LC-MS systems compromises instrument performance and is thus not a practical solution.

Our observations (Fig. 4), and that of others ^28, 90^, reveals a marked difference in the propensity of PTM neutral loss in both HCD and CID between sY- and pY-peptides. We thus hypothesised that we could exploit this feature to develop a low energy neutral loss EThcD triggering approach that could discriminate near-isobaric PTMs and enable sY-peptide identification from complex mixtures containing pY peptides. To test this, we quantified HCD precursor ion neutral loss at 10% NCE for sY- and pY-peptides, quantifying -80 Da (SO_3_/HPO_3_) and -98 Da (H_2_SO_4_/H_3_PO_4_) mass shifts (Fig. 5). For each peptide spectrum match (Fig. 4), the relative abundance of the precursor or neutral loss precursor ions were calculated as a percentage of total product ion current (Fig. 5). The predominant 10% NCE HCD product ion observed across our sY-peptide panel equated to loss of 80 Da (sulfonate) from the precursor, accounting for 20-100% of the MS/MS ion current (median = ∼85%). Little to no peptide backbone cleavage was observed, in agreement with the finding that no peptides were identified with the search engine under this condition (Fig. 4). In contrast, pY-peptide exhibited minimal loss of either 80 or 98 Da (<1%) under the same conditions (Fig. 5) (Supp. Fig 1).

**Figure 5:**
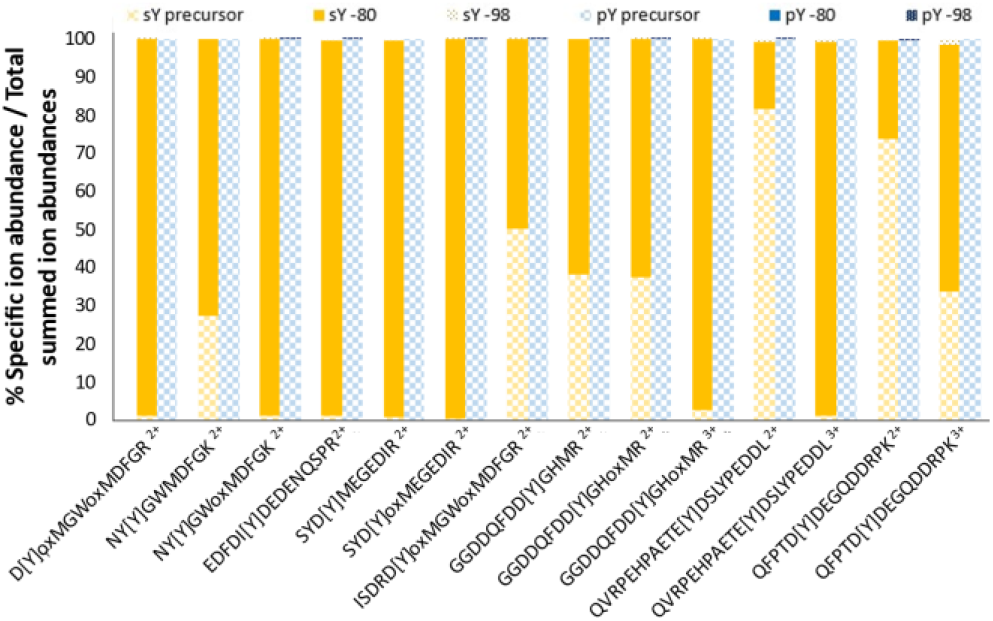
Neutral loss propensity of sY- versus pY- peptides at 10% NCE HCD fragmentation. The most intense PSM (determined from the mgf file) for each sY and pY peptide at different charges (including oxidised methionine variants) were selected for quantitation (14 peptide ions). The relative abundance (%) of the precursor ion (check), precursor ion -80 Da (block), or precursor ion -98 Da (dotted) was calculated as a proportion of the summed intensity of all ions in the MS2 spectrum. sY-containing peptides in yellow; pY peptides in blue.

Interestingly, while the degree of neutral loss for sY peptide ions appeared to be dictated by the ratio between the number of basic residues and the charge state, the trend was the opposite to that observed for phosphopeptides.^91^ Where the charge state was greater than the number of basic residues (H/K/R), contributing to a ‘mobile proton environment’ (MPE), we observed near complete neutral loss of 80 Da. However, the degree of precursor ion neutral loss was comparatively reduced when the charge state was smaller than the number of basic sites (−80 Da neutral loss: QFPTD[Y]DEGQDDRPK: 2+ ∼20%, 3+ ∼65% and QVRPEHPAETE[Y]DSLYPEDDL: 2+ ∼15%, 3+ ∼95%). These data suggest that the charge-directed fragmentation mechanisms that appear to drive phosphopeptide neutral loss^91, 92^ are not directly applicable to sulfopeptides, whose fragmentation (propensity for neutral loss) is also likely to be affected by the differences in electronegative and hydrogen bonding capabilities of the sulfonate moiety.

Having demonstrated the ability to distinguish sY- and pY-peptides based on HCD precursor ion neutral loss at 10% NCE, we next sought to implement this for sY-peptide identification using a neutral loss triggering approach in a mixture. Given optimal peptide identification across all charge states with HCD (32% NCE) (Fig. 4), we used the 80 Da precursor ion loss at 10% NCE HCD to trigger 32% NCE HCD on the same precursor to permit peptide identification, analysing a 1:1 ratio of our sY- and pY-peptide panel. Using this approach, we were able to positively identify all of the sY-peptides, and did not identify any of the pY-containing peptides, confirming the utility of this low energy HCD triggering strategy. To our knowledge, this is the first reported case of utilising low collision energy HCD fragmentation to efficiently distinguish sY- and pY-peptides by MS in a single run.

### Application of our HTP sulfopeptide analytical pipeline to identify the secreted ‘sulfome’ of HEK-293 cells

sY-proteins are generated in the Golgi compartment, and are predominantly destined for secretion or cell membrane localisation: of the 33 validated human sY-proteins in UniProt, 19 are secreted and 14 are membrane bound.^7^ In order to test the capabilities of our optimised workflow for sTyr, we evaluated the secretome of the adherent HEK-293 model cell system (Figure 6). After an 18 h incubation in serum-free medium, the HEK-293 cell secretome was collected, purified and prepared using Strataclean resin and an SP3-based trypsin digestion protocol (adapted from ^93^) and treated with protein phosphatase.

**Figure 6:**
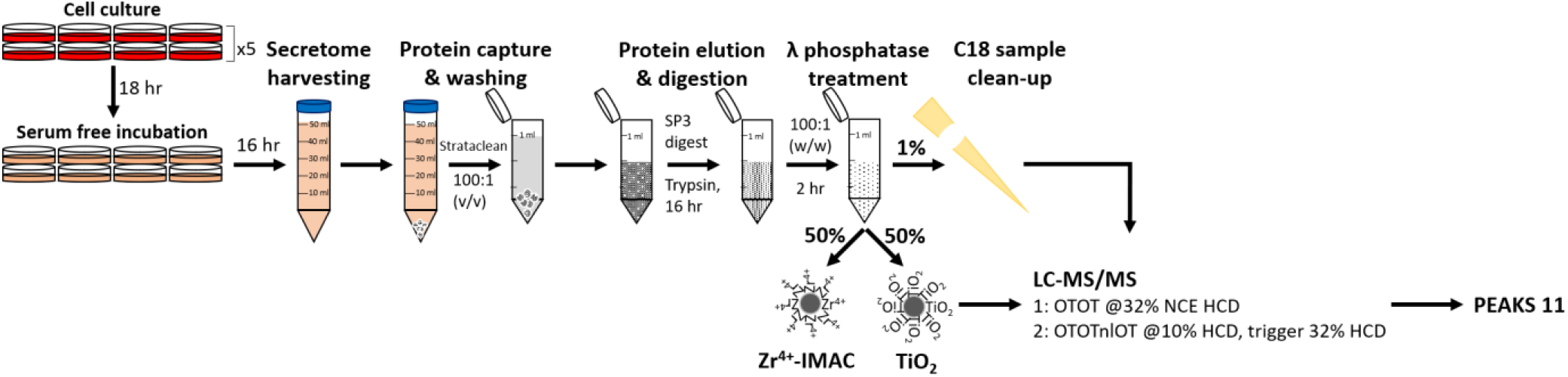
Workflow for the identification of sulfated peptides from a HEK-293 cell secretome. Following incubation in serum-free media, the secretome was harvested, and proteins captured with Strataclean beads. Eluted proteins were subject to SP3-based trypsin proteolysis and treatment with λ phosphatase. 1% was subjected to C18 sample clean-up prior to LC-MS/MS analysis. The remainder was subject to enrichment using either the optimised Zr^4+^-IMAC or TiO_2_ protocols. All samples were analysed using i) high resolution (Orbitrap, OT) data-dependent acquisition (DDA) with 32% NCE HCD, or ii) the neutral loss triggering strategy where loss of 80 Da at 10% NCE HCD invoked precursor fragmentation with 32% NE HCD. Data were analysed using PEAKS 11.

Given differences in pY and sY peptide enrichment between Zr^4+^-IMAC and TiO_2_ protocols, we elected to compare both optimised enrichment protocols to investigate the cellular secretome. However, initial analysis of enriched material from agarose coated (Purecube) Zr^4+^-IMAC resin using a standard data-dependent acquisition (DDA) pipeline revealed extensive binding of non-modified peptides. We thus employed MagReSyn® Zr-IMAC HP (ReSyn Biosciences), which exhibits substantially reduced non-specific peptide binding.^94^

To allow us to evaluate (i) the efficiency of sY- versus pY-peptide enrichment using the TiO_2_ and MagReSyn® Zr^4+^-MAC HP resins, and (ii) the neutral loss triggered-strategy for sulfopeptide identification, we acquired MS/MS data using a DDA pipeline alongside our neutral-loss triggered approach (Figures 6, 7).

**Figure 7:**
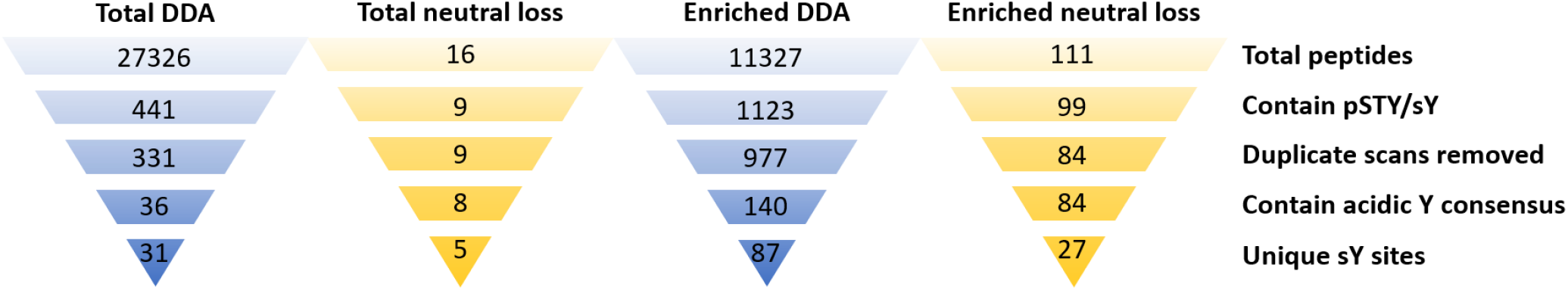
Evaluation of sulfopeptide enrichment and neutral loss triggering data acquisition for sulfopeptide identification. Secretome samples were analysed using either a data-dependent acquisition (DDA) strategy (32% NCE HCD) or the 10% HCD neutral loss triggered strategy for HCD acquisition (32% NCE), with or without enrichment using TiO_2_/ Zr^4+^-IMAC enrichment (Figure 6). Listed are the total number of peptides identified, those containing either pSTY or sY before or after removal of duplicate scan PSMs. Peptide lists were subsequently filtered for those where the site of modifications was in an acidic consensus (with D/E at the +1 or -1 position relative to Y), and then for unique sY sites.

To confirm effective purification of the secretome, 1% of the sample was subject to DDA analysis using HCD (NCE 32%). Performing an initial open PTM search (PEAKs PTM) we identified a high degree of Met oxidation (∼5,900 peptide spectral matches (PSMs)) and Asn/Asp deamidation (∼5,500 PSMs). Met oxidation and Asp/Asn deamidation, as well as Ser/Thr/Tyr phosphorylation and Tyr sulfation were thus included as variable modifications in subsequent searches. From the total secretome DDA analysis, we identified 27,326 peptides from 2,695 proteins at a 1% FDR (Figure 7). Gene Ontology (GO) analysis with DAVID (Supp. Table 2) revealed that 36% of the identified proteins were defined as being localised to extracellular exosomes (Benjamini-Hochberg corrected p-value <1.6e^-163^), 29% as membrane-bound (Benjamini-Hochberg corrected p-value = 6.49e^-56^) and 44% as cytosolic (Benjamini-Hochberg corrected p-value = 7.72e^-101^). Of these ∼27k peptides, 441 (<2%) were annotated as containing either pS/T/Y or sY (330 and 141 sites respectively) (Figure 7). A substantive proportion (∼25%) of these annotations were from duplicate scan numbers where the same scan generated PSMs annotated as containing either pSTY or sY, or differences in deamidation status of Asn/Asp. Manually filtering this list, retaining the highest scoring scan number unique PSMs yielded 331 identifications. Enzymatic deposition by TPST1/2 is known to occur on Tyr residues within an acidic consensus (with Asp or Glu localized at either the +1 or -1 position), therefore we further filtered these identifications based on an acidic consensus, revealing 36 peptides, 31 of which were unique sites of Tyr sulfation (Figure 7).

A total of 11,327 peptides were identified following DDA analysis of both enriched samples. No substantive differences in terms of isoelectric point or *m/z* distribution were observed for those PSMs identified from the total or the enriched secretome samples (Supp. Fig. 2). Ten percent of the enriched samples (1,123) were annotated as phosphorylated and/or sulfated (898 and 304 respectively). While markedly lower than the level of enrichment typically seen from a cell extract using standard phosphopeptide enrichment strategies^58, 95-97^ (usually ∼85-90% using TiO_2_ in our hands), this equates to over a 6-fold increase in the identification of modified peptides from the secretome sample, and a ∼2.2-fold increase in the peptides annotated as being sulfated. Removal of duplicate scans (977 identifications) and filtering for an acidic consensus left some 140 peptides with 87 sites (Figure 7).

Performing our neutral loss triggered MS acquisition method with the same sample significantly reduced the number of peptides identified in both the total (0.06%) and enriched (1.0%) samples. The numbers of peptides identified in the enriched DDA experiment compared to the enriched neutral loss analysis indicates that the enrichment strategy alone is insufficient for enhanced sulfopeptide identification (in agreement with our synthetic peptide analysis, Figure 3), but that enrichment does serve to improve the sensitivity of the MS acquisition.

Of the two different enrichment strategies, the sample prepared using Zr^4+^-IMAC HP resin triggered the greatest number of MS/MS spectra (demonstrating higher likely enrichment of sY-peptides) following 10% NCE HCD, yielding 977 MS2 spectra; the TiO_2_-enriched sample resulted in 754 triggering events. Despite >1700 total triggering events, only 327 spectra were matched to a peptide sequence (∼19%) suggesting issues associated with fragmentation, and/or ion intensity that compromised identification. Surprisingly, of these 327 PSMs, 319 (∼98%) were from the TiO_2_ sample, with only 8 (∼2%) spectra from Zr^4+^-IMAC HP resin. We are currently at a loss to explain this marked difference in the ratio of triggering events to peptide identifications, noting that the TiO_2_-enriched samples generally yielded ions of lower average intensity with a lower charge state distribution (Supp. Fig. 3). In terms of total peptide identifications – this equates to 111 for TiO_2_ of which 62 were annotated as sY-containing (54% enrichment efficiency), and 7 for Zr^4+^-IMAC of which 6 were annotated as sulfated (86% enrichment efficiency). Applying the acidic-Tyr filter and concatenating for the highest scoring PSM per scan for our enriched neutral loss triggered dataset (Figure 7), we identified 84 modified peptides, all of which contained a Tyr residue within an acidic motif. The resultant 27 unique sites of modification mapped to 23 proteins (Table 1). To note, only 5 of these sites (from 4 proteins) were found in the unenriched DDA experiment.

**Table 1:**
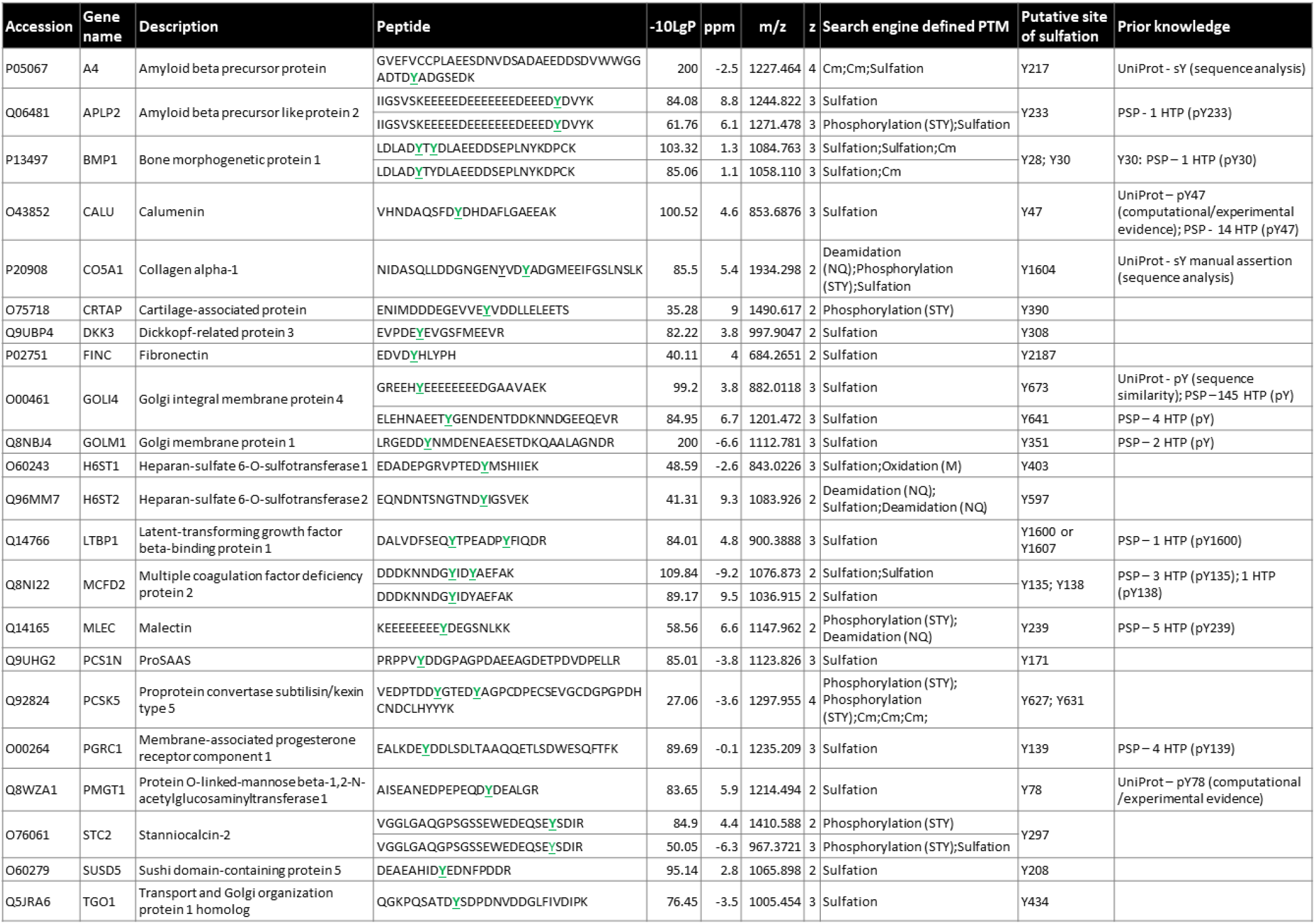
Confidently identified sY-peptides from a HEK293 cellular secretome. Detailed are the accession number, gene name, protein description, peptide sequence, -10LogP score, Δppm from theoretical *m/z*, the observed *m/z*, charge state (z) and identified PTMs as determined by the search engine. Listed is a concatenated version of all modified peptide identifications, maintaining only the highest - 10LogP score peptide for a unique sY-site. Predicted (acidic) sY-site is **<u>highlighted</u>**. Cm – carbamidomethylation. All predicted sY sites were searched for prior annotation as being either phosphorylated or sulfated in UniProt and the PhosphositePlus (PSP) database [accessed July 2023]; HTP – details the number of times phosphorylation was reported in PSP in a high throughput study.

Considering that our neutral loss triggering MS acquisition method can efficiently discriminate sY-from pY peptides, we attempted to utilise confidence scores (−10LogP) and Δppm to distinguish between pSTY/sY of these duplicated scans (Supp.Fig 4). The majority of PSMs were annotated as being sulfated, with or without N/Q deamidation. Although Δppm was generally lower for those PSMs with a higher -Log10P value, this was not consistent. PSMs annotated as deamidated generally had both a higher Δppm and were lower scoring. We did not observe any score or ppm-related features that could confidently differentiate assigned PSMs as being either phosphorylated or sulfated. We thus concatenated the data, removing the lower scoring duplicate PSM to retain a single identification per scan – this yielded a final dataset of 23 different peptides and 30 pSTY/sY annotations from the HEK-293 secretome (Table 1).

We next compared our dataset of 27 modified peptides with the human Tyr ‘Sulfome’ in UniProt. Interestingly, only Y217 on Amyloid beta precursor protein (A4, P05067) and Y1604 on collagen alpha-1 (CO5A1, P20908) have been previously identified as sites of sulfation. While other sites of sY have been reported in both of these proteins, these were not identified in our data. A number of these, including Y262 on A4 is situated in extremely acidic regions which likely do not lend themselves to trypsin-based identification.

Six of the 27 peptides contain multiple sites of (+80 Da) modification, being annotated as either doubly sulfated, doubly phosphorylated, or a mixture of the two. Focussing in the first instance on a doubly ‘phosphorylated’ peptide from PCSK5 (Proprotein convertase subtilisin/kexin type 5), two of the five Tyr residues have an Asp at -1, and are localised within a highly acidic region (the others being clustered at the peptide C-terminus). It is also worth noting that the tandem MS data that generated these identifications was triggered based on neutral loss of 80 Da at 10% NCE HCD, which we have shown does not induce phosphate loss (Fig. 5, Supp. Fig. 1). Like the mixed PTM containing peptides (sY and pS/T/Y) identified from APLP2 (amyloid β precursor like protein 2), CO5A1 (collagen α-1) and STC2 (stanniocalcin-2), it is therefore likely that (at least) one of these sites is in fact sulfated; we assume that neutral loss from the sulfated residue under lower energy conditions triggered MS/MS data acquisition and identification of a peptide that is **also** phosphorylated. Likewise, the singly modified site on STC2 (which is embedded in an EYxD motif) is likely also sulfated. To investigate this further, we interrogated the 10% HCD scans for these precursors, determining that the MS2 spectra from both APLP2 and STC2 contained a neutral loss peak equivalent to a single sulfated residue (−80 Da), while spectra for the secreted metalloproteinase BMP1, MCFD2, CO5A1, PCSK5 contain losses equivalent to (80 and) 160 Da, suggestive of two sites of tyrosine sulfation. This agrees with information in UniProt that identifies Y1601 on CO5A1 as an additional site of sulfation, strongly indicative of mis-identification of sulfation sites as phosphosites by the search engine.

Given our synthetic peptide panel data (Fig. 4), an unexpected finding from our final secretome-derived (non-unique) dataset was the large number of ions (61 out of the 84 peptides i.e. >60%) that were observed with a charge state ≥3. The sulfation-induced reduction in charge state observed for the panel meant that we had expected a substantive proportion of those peptides observed using our sulfoproteome pipeline (Figure 6) to be doubly protonated. In fact, while this was true across all peptides in the DDA dataset from the enriched secretome, this was not the case for the NL triggered dataset, with the majority of peptide ions appearing as 3+ species (Supp. Fig. 5A). We believe that this is because the peptides identified with the NL triggering approach are much larger (and thus of higher mass) compared to those from the enriched DDA set (Supp. Fig. 5B, C) or our peptide panel; this can be explained by virtue of the fact that the sY-containing (NL-triggering) peptides from the secretome contained a substantially higher proportion of acidic residues (9.0 D/E residues on average) compared with those peptides in the total protein DDA set (average of 1.7 D/E residues) (Supp. Fig. 5D), and consequently a lower relative proportion of K/R residues.

To validate the secretome data obtained using our pipeline for identification of Tyr sulfation, we expressed and affinity precipitated the novel sulfated substrates H6ST1 and H6ST2 (Golgi-localized Heparan-sulfate 6-O-sulfotransferase 1/2) from HEK-293T cells, and subjected both proteins to *in vitro* PAPS-dependent sulfation with recombinant TPST1/2. We also immunoprecipitated the related isoform H6ST3, which contains an analogous Tyr residue within the acid consensus sequence in its C-terminal region. After enzymatic reaction, proteins were digested with trypsin and analysed by LC-MS/MS using both DDA and our neutral loss triggered method. As well as confirming TPST1/2-dependent sulfation of H6ST1 and H6ST2 at the same sites on tryptic peptides observed from our HTP study (sTyr403 and sTyr597 respectively, Supp. Tables 3, 4), we also show that the related protein, H6ST3 can also be Tyr sulfated at sTyr285 and sTyr464, Supp. Tables 3, 4). Of particular interest, sTyr403, sTyr597 and sTyr 464 lie in a conserved acidic motif in the C-terminal region of H6ST1-3 outside of the catalytic domain (Supp. Fig. 6). Not only does this analysis validate the exploratory potential of our discovery pipeline for sTyr detection, but it also demonstrates how this information can be extrapolated to predict additional sites of modification in closely-related proteins such as H6ST3. Interestingly, only one of the two TPST1/2-dependent sulfation sites on HS6ST3 contained an acidic residue in close proximity (sTyr464 – [ED**<u>Y</u>**X]), suggesting that an acidic motif around the site of sulfation may not be an absolute requirement for TSPT1/2 substrates.

As well as validating our HTP sulfomics pipeline, our enzymatic assays also revealed additional novel TPST1/2-dependent sulfation sites on proteins that co-immunoprecipitated with H6ST2 and/or H6ST3: Gem-associated protein 5 (GEMIN5; sTyr992), tubulin α-1A/B/C chain (TUBA1A/TUBA1B/TUBA1C; sTyr161, sTyr432), Tubulin β chain (TUBB; sTyr[50 or 51], sTyr340), insulin receptor substrate 4 (IRS4; sTyr921) and heterogeneous nuclear ribonucleoprotein H (HNRNPH1; sTyr266). Gemin5 is thought to reside in the nucleoplasm and in specific nuclear bodies (Gemini of Cajal Bodies), as well as the cytoplasm, and has not previously been shown to be sulfated (or modified on Tyr992). Likewise, none of the other TSPT1/2 substrates have previously been reported to be sulfated. However, the *in vitro* sulfated Tyr sites identified on these proteins have previously been reported as being Tyr phosphorylated in PhosphoSitePlus in HTP proteomics screens, with IRS4 Tyr921 described as a putative substrate for the Tyr protein kinases Fer and IGFR1. Finally, we also identified what we believe to be the first auto-Tyr sulfation site on TPST1 (likely at Tyr326), which might itself be relevant to cellular TPST1/2 regulation or enzyme:substrate interactions, similar to the signaling paradigm in which tyrosine kinases are themselves controlled by tyrosine phosphorylation.

Together, these data provide strong evidence that our sulfomics pipeline is capable of defining novel human sulfopeptides in an HTP manner, and suggest Tyr sulfation cross-talk as a potential regulatory modification between Golgi co-localized human heparan sulfate sulfotransferases and TPSTs.

## Conclusions

In this study, we investigated several strategies for the analytical discrimination of sulfopeptides from phosphopeptides by employing, to our knowledge, the largest panel of synthetic peptides based on tryptic human peptides derived from known sites of sY-modifications from cellular proteins. In developing a workflow specifically for HTP ‘sulfomics’ we characterised a number of sY-peptide discriminatory features that include RT shifts, susceptibility (or lack thereof) to treatment with protein phosphatase, conditions for sY-peptide enrichment and peptide fragmentation. We developed, and implemented the first HTP based ‘sulfomics’ pipeline and demonstrated its utility using a HEK-293 cellular secretome, where we identified a total of 21 novel (experimentally-determined) sY-containing proteins and 28 sY-sites, which expands the known human ‘sulfome’ by ∼70%. In the process of validating H6ST1 and H6ST2 (and HS6ST3) as *in vitro* targets of the sulfotransferases TSPT1/2, we also identified additional *in vitro* substrates – GEMIN5, TUBA1, TUBB, IRS4 and HNRNPH1, which are themselves part of the known H6ST2/3 interactome. Interestingly, while the site on GEMIN5 has not previously been identified as being modified, there is extensive HTP evidence of phosphorylation of the other TSPT1/2 substrates. It thus remains to be seen whether i) these sites can be **both** sulfated and phosphorylated *in vivo*, ii) our identification of these specific sulfation sites *in vitro* is biased by the assay conditions, or iii) the phosphorylation site identifications reported in PhosphoSite Plus are actually mis-representations of a Tyr sulfation event. Since Tyr sulfation is believed to be irreversible, our findings raise the possibility that false identification of sTyr as pTyr, may be meaningful for several proteins, and urgently needs to be clarified for multiple proteins. In addition, since sTyr could act as a Golgi-based signal that competes with, or prevents, Tyr phosphorylation further along the secretory pathway, we are in the process of evaluating this combinatorial phenomenon further. Regardless, the identification of both glycan and protein sulfotransferases as sY-containing proteins potentially opens up a new research area for understanding the regulation and substrate targeting of these enzymes, similar in many ways to the phosphoregulatory paradigms established for protein kinases.^98^

While Fe^3+^/Ga^3+^ IMAC has previously been used for sY peptide enrichment ^26,27^, we were unable to validate these findings using our peptide library. However, we demonstrate the utility of Zr^4+^-IMAC and TiO_2_ in acetic acid-based solutions for the semi-specific enrichment of sY-peptides in a complex (phospho) peptide mixture. Noting the differential enrichment of the relatively acidic non-modified peptides from our library (average pI ∼4.5), compared with unmodified peptides from BSA/casein (average pI ∼5.7), our evidence suggests that the acidic consensus (thought to be required for TPST1/2 directed sulfation) also contributes to the efficiency of sY-peptide enrichment. This is also reflected in the complex secretome analysis where the neutral loss triggered enriched peptides (sY-containing) identified were of much greater acidity than the total secretome DDA identifications (pI ∼4.6 versus ∼6.2).

In undertaking comprehensive MS analysis of tryptic sulfopeptides using CID, HCD, EThcD, ETciD and UVPD fragmentation regimes, we were able to quantify the generation of site-localising product ions and compare them with equivalent synthetic phosphopeptides. Taken together, our data indicates that specific sulfosite localisation is poor, irrespective of fragmentation regime or conditions used. Our data with the commercially available UVPD configuration did not prove as promising as anticipated, based on previous work.^43,44,39^ A number of factors likely contribute to this: difference in UVPD wavelength, energy and/or laser frequency, the use of positive rather than negative ion mode, and the fact that many of the studies reported to date use a handful of exemplar peptides. EThcD appeared to be the best overall for site localisation, but was inconsistent in its ability to identify sulfopeptides, in part because of the reduction in charge state. During the course of our studies we determined that low energy (10% NCE) HCD could serve to differentiate sulfopeptides from phosphopeptides, based on the lability of the sulfate group. We thus made use of the single feature that caused the greatest analytical challenge in terms of localisation to allow sY discrimination ‘on the fly’; a low energy (10% NCE) HCD neutral loss-triggering approach to define sulfopeptide-containing scans for subsequent identification. Our initial thinking based on the reduced ionisation efficiency of sulfopeptides was that application of EThcD to global ‘sulfome’ analysis would be compromised. However, our secretome-derived dataset suggests that the average increase in length and charge state of sulfopeptides may make a neutral-loss triggered EThcD strategy feasible for HTP sulfosite identification. However, this strategy will need to be evaluated for throughput given the additional time requirements of EThcD over HCD, and the necessity given that i) most peptides only contain a single Tyr residue and ii) the strong requirements for an acidic consensus for TSPT1/2 activity.

Thus, while the pipeline described herein facilitates the identification of sY-peptides from complex mixtures, unambiguously pinpointing the precise site Tyr of modification remains challenging. Specifically, the electronegativity of sY and issues associated with site localisation in positive ion mode MS (given current fragmentation strategies) continue to compromise definitive site determination, unless peptides only contain a single Tyr residue.

Recent reports have commented on the inherent metal ion-binding affinity of sY that might be exploited for localisation (^99, 28^ & ^90^), with one report identifying metal adducted species as dominant when using standard positive ion mode LC-MS/MS methods (specifically Na+/K+, ^64^). As such, we performed open PTM searches on our datasets in an attempt to identify some of the missing triggered peptides (86% of triggers). However, we failed to observe these species in our investigations. No additional sY-containing proteins were identified with the open PTM search, although differentially modified species of already identified sY peptides were observed. We presume this is a direct result of how PEAKs PTM performs its searches, requiring a protein to be identified by a first round database search with pre-defined search parameters, prior to performing a 2^nd^ round mass shift search on a concatenated database containing only these identified proteins.^100^ Since our neutral loss triggering method drastically reduces the number of peptides and proteins identified, this compromises the ability for PEAKS PTM to identify additional sulfated peptides in the absence of defining specific additional modifications. In summary, we believe that the analytical developments reported in this paper will serve as the basis of a new pipeline for HTP sulfome analysis, providing a resource to the community to better define the extent and roles of protein sulfation from biological samples.

## ASSOCIATED CONTENT

### Supporting Information

Supplementary Figures and Supplementary Table 3 – PDF Supplementary Tables 1, 2 and 4 - .xlsx

All mass spectrometry data has been deposited at the ProteomeXchange Consortium (http://proteomecentral.proteo-mexchange.org) via the PRIDE partner repository with the dataset identifiers: Project Accession - PXD043713 and PXD043723.

## AUTHOR INFORMATION

### Author Contributions

LAD: performed all sample preparation, enrichment and mass spectrometry analysis. Contributed to design of experiments and manuscript writing. DPB: performed biochemical analyses – phosphatase experiments and sulfation assays. SP & EM: contributed to MS/MS fragmentation data analysis. PJB: supported mass spectrometry methods development. ARJ: contributed to MS/MS fragmentation data analysis. PAE: contributed to design of experiments and manuscript writing. CEE: contributed to design of experiments, analysis of data and manuscript writing.

The manuscript was written by LD and CEE with contributions from PAE, DPB and ARJ. All authors have given approval to the final version of the manuscript.

## ACKNOWLEDGMENT

This work is supported by from funding from the Biotechnology and Biosciences Research Council (BBSRC; BB/S018514/1, BB/M012557/1, BB/S017054/1 and BB/R000182/1). We thank Prof David Fernig (University of Liverpool) for useful discussions.

## Notes

### Competing Interest Statement

The authors have declared no competing interest.

