## Supplemental Figures and Supp Table 3 for "A bespoke analytical workflow for the confident identification of sulfopeptides and their discrimination from phosphopeptides"

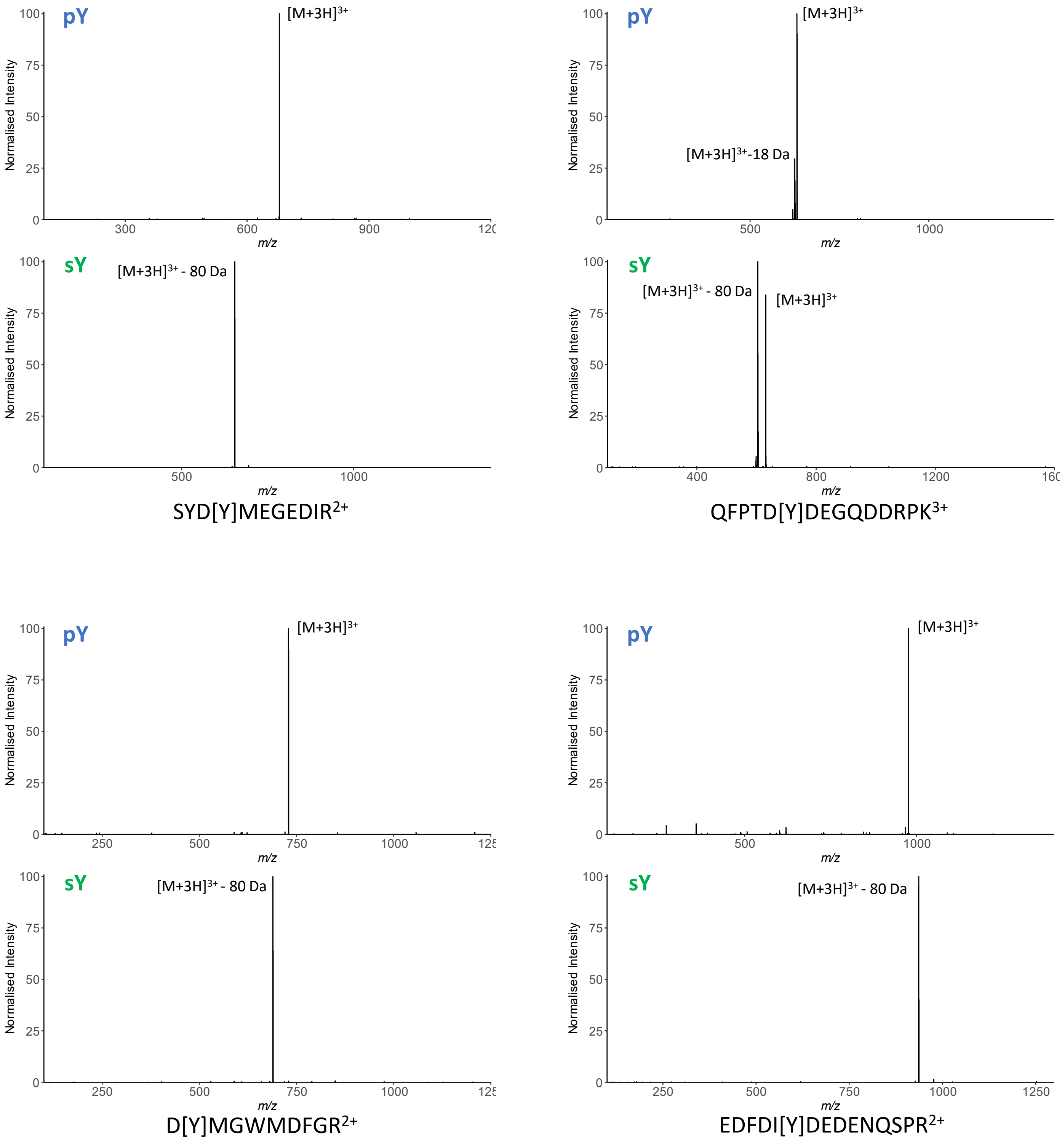


**Supp. Fig. 1: Neutral loss propensity of sY versus pY-containing peptides at 10% HCD.** MS2 spectra of identical peptides containing either sY or pY were subjected to 10% HCD fragmentation. Sequence and charge state of exemplar peptides are displayed, the identification of the m/z peaks are annotated. Spectra redrawn using a custom R script from the .mgf file.

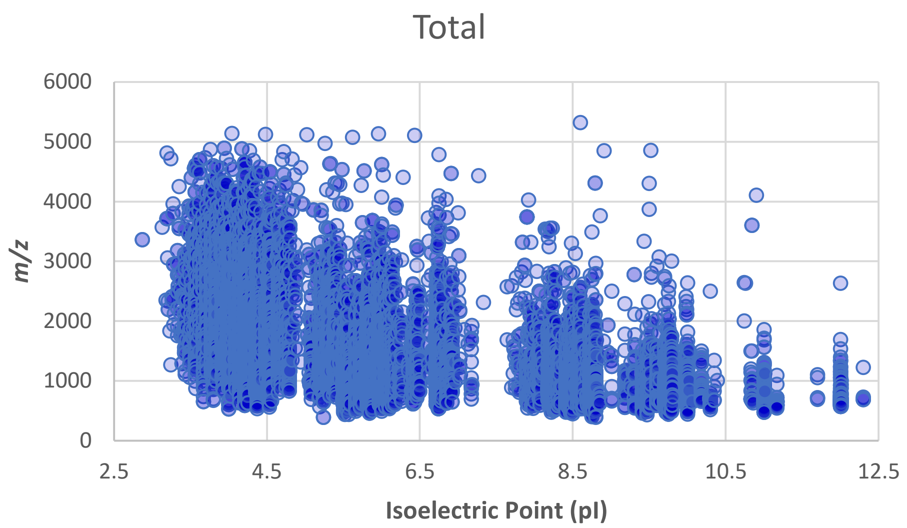


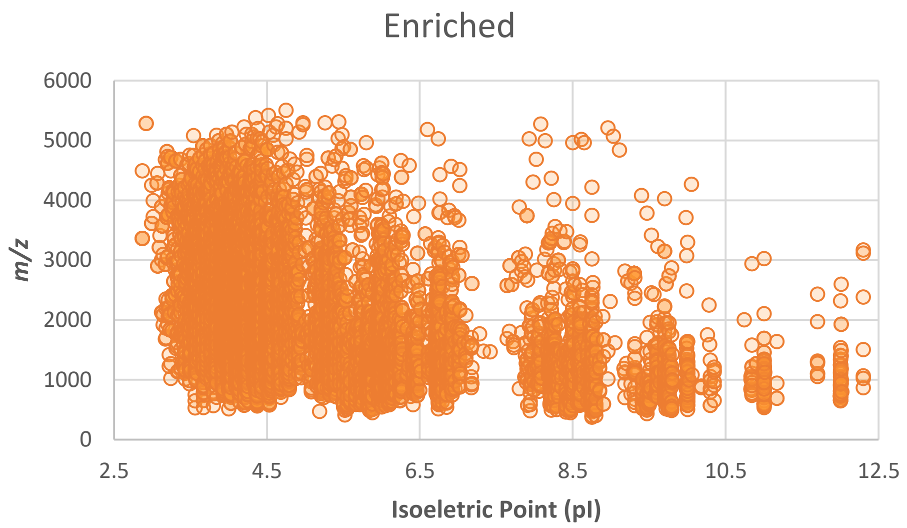


**Supp. Fig. 2: Isoelectric point versus *m/z* distribution of the peptides identified from either the total DDA unenriched HEK-293 secretome sample (top; blue), or following enrichment with $TiO_2$ or $Zr^{4+}$-IMAC (bottom; orange).** m/z data was extracted from PEAKs11 analysis. Identified peptide sequences were input into the Expasy compute_pI tool to determine pI (https://web.expasy.org/compute_pi/) [accessed June 2023].

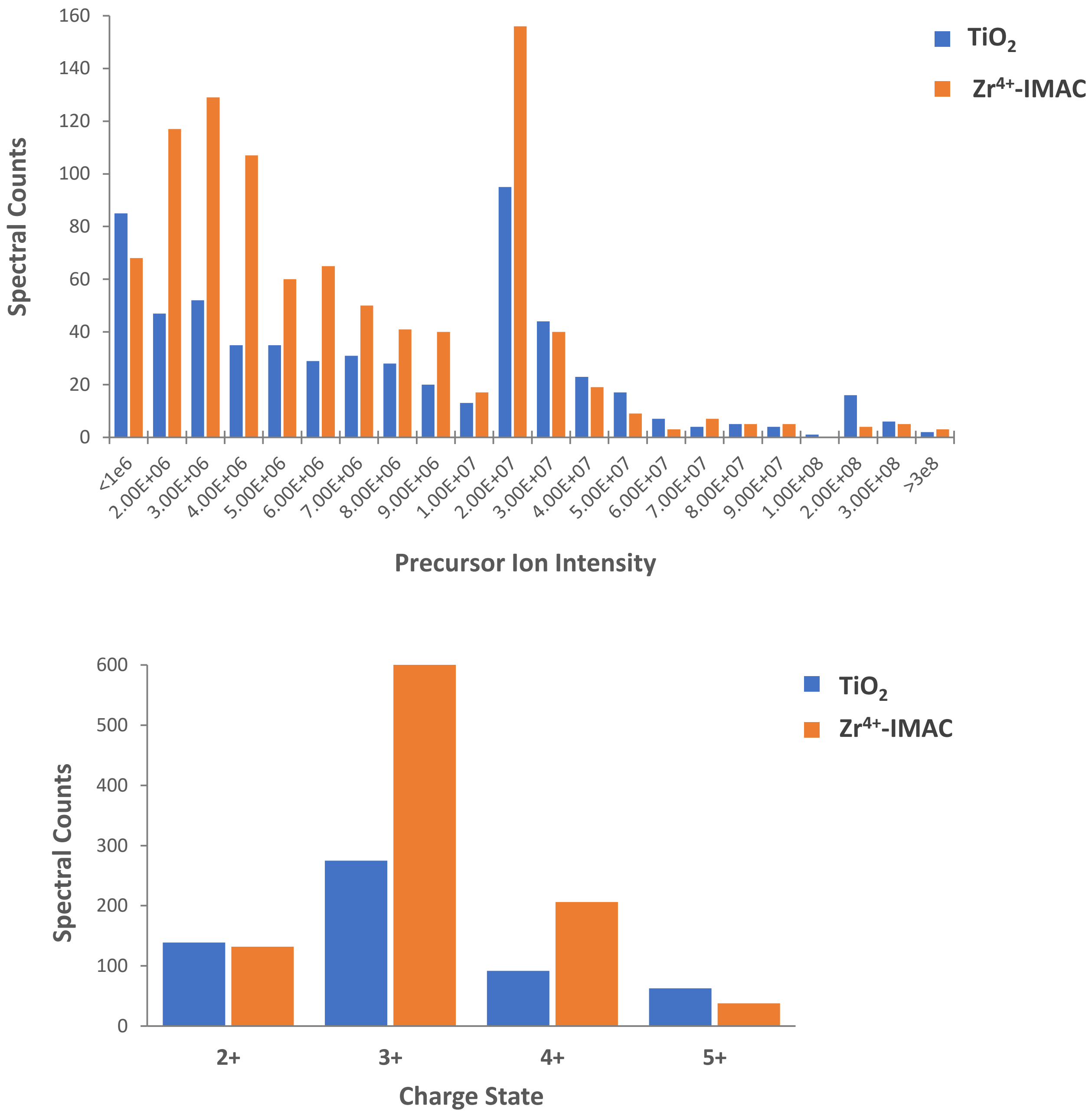


**Supp. Fig. 3: Characteristics of precursor ions that triggered 32% NCE HCD following 10% NCE HCD -80 Da neutral loss.** Precursor ion intensity distribution (top) or charge states (bottom) for either the $TiO_2$ (blue) or $Zr^{4+}$-IMAC (orange) enriched HEK-293 cell secretome samples. Data was extracted from .mgf files where the MSconvert "HCD energy = 32" filter was set to specifically identify triggered scans.

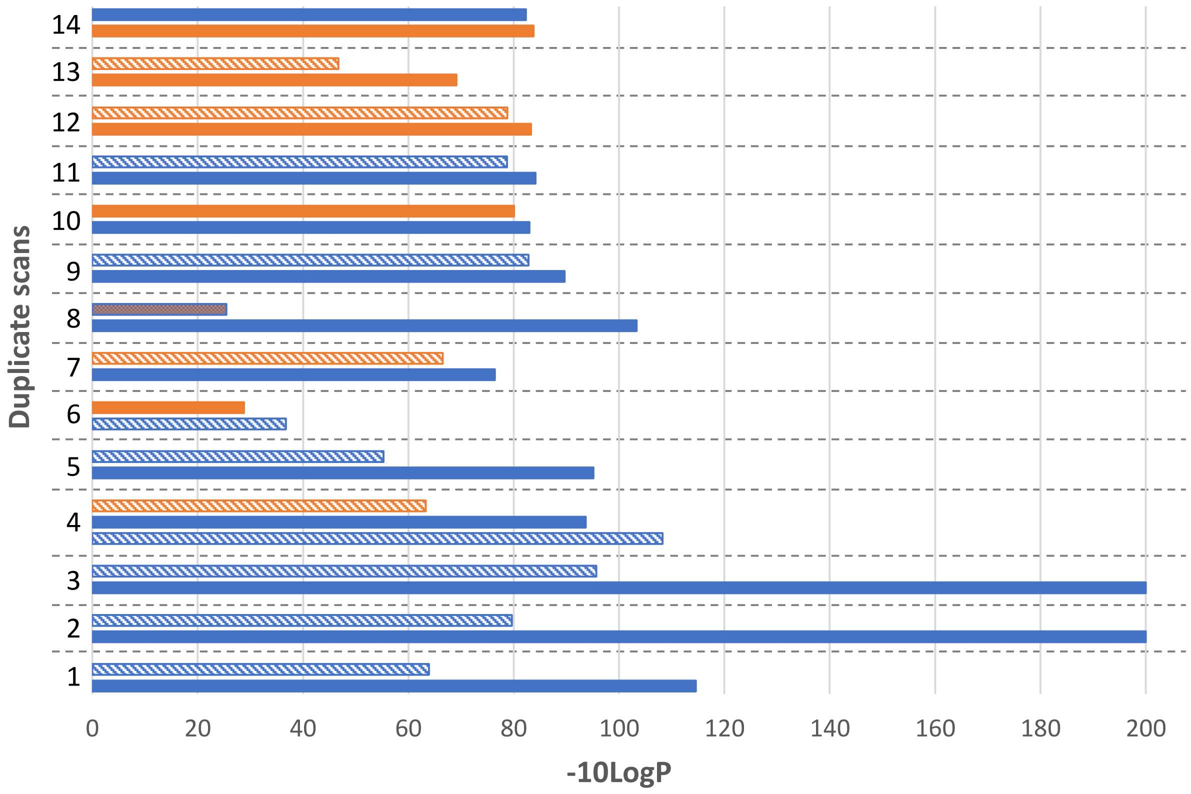


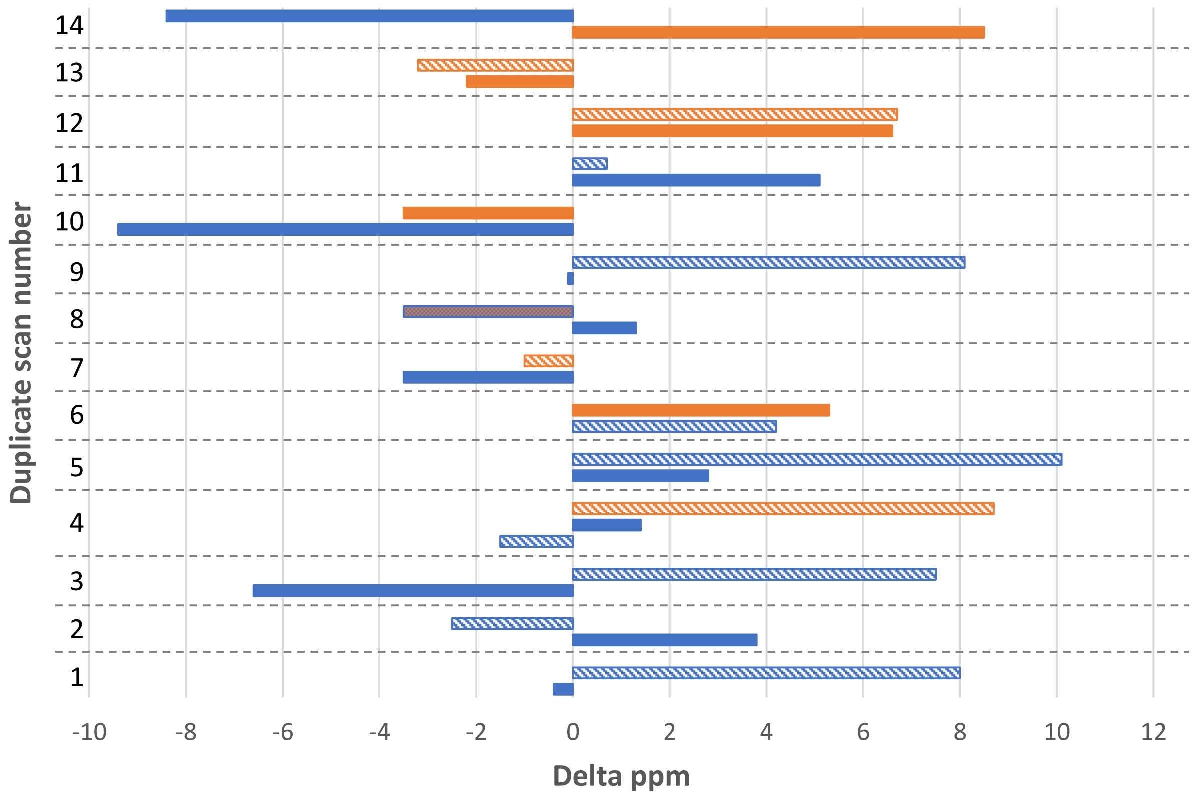


**Supp. Fig. 4: Distinguishing duplicated scan PSMs.** -10LogP value (top) and Δppm (bottom) for 14 representative duplicate scans from the neutral loss triggered enriched dataset with different PTMs assigned to the same peptide sequence. Sulfation - blue; phosphorylation – orange; deamidation – check boxes. Better score often, but does not always, correlate with a lower Δppm.

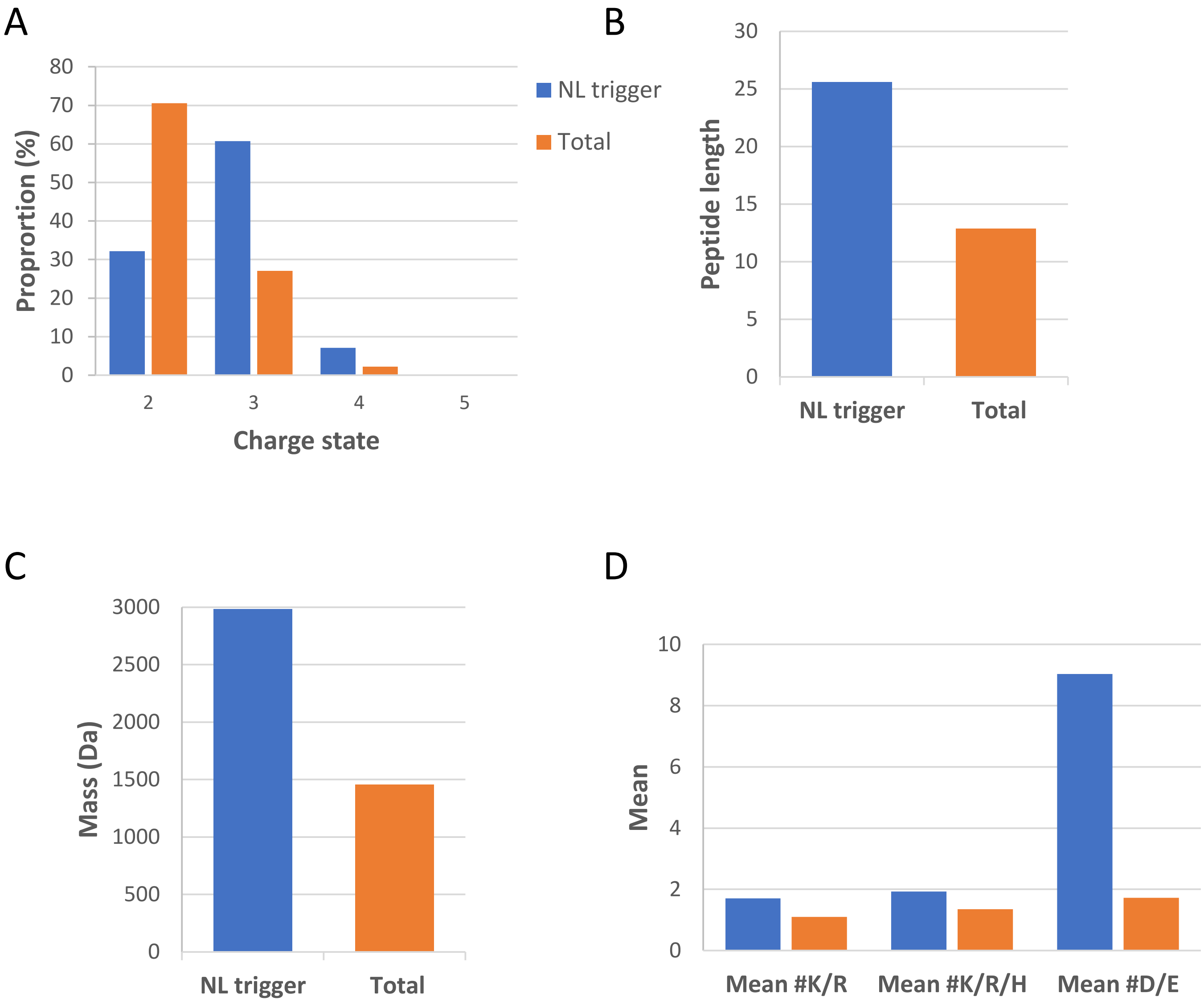


**Supp. Fig. 5: Properties of peptides from enriched HEK293 secretome.** All non-duplicated PSMs from either the neutral loss triggered enriched (blue) or the total unenriched protein DDA (orange) analyses were interrogated for (A) charge state; (B) peptide length; (C) average mass; (D) average (mean) number of K/R/H or D/E residues per peptide. All data was extracted from the PEAKs11 output.

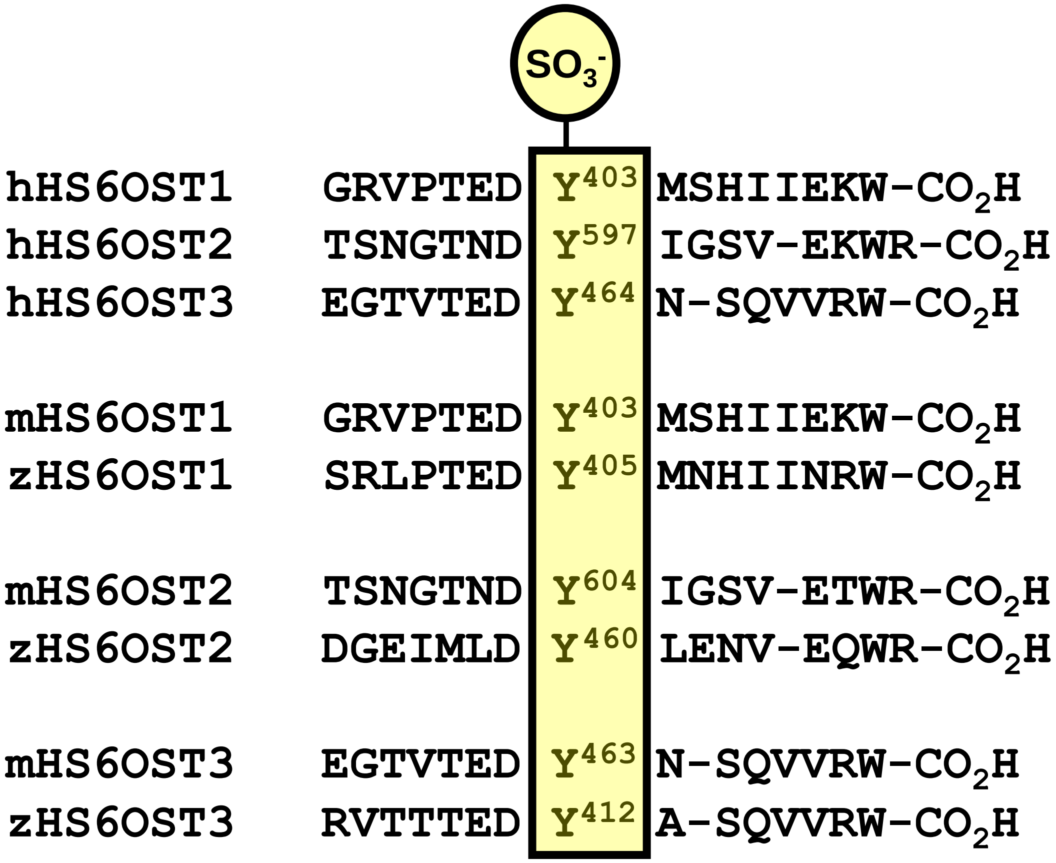


**Supp. Fig. 6:** Conservation (shaded) of a validated sulfated Tyr residue in an acidic C-terminal motif in human (h) Heparan Sulfate 6OST1, 2 and 3. The aligned sequence of the equivalent C-terminal region from mouse (m) and zebrafish (z) is also shown.

**Supp. Table 3:** ***In vitro*** **TPST1/2 sulfation assay.** H6ST1, H6ST2 or H6ST3 were overexpressed in HEK-293 cells, immunoprecipitated and subjected to *in vitro* sulfation with TPST1/2. Samples were digested with trypsin and subjected LC-MS/MS analysis, using either data-dependent acquisition (DDA) or the low energy (10% NCD HCD) neutral loss (NL) triggering strategy. Detailed are the conditions for the assay, identified proteins of interest, protein sequence coverage (%) and Mascot score, sulfopeptide sequence, region in the protein and peptide score. Putative site of sulfation in the peptide sequence is underlined. No modified peptides were observed in the absence of TPST1/2.

| Condition | Gene name | Protein name | % Coverage | Mascot Score | Sulfopeptide Sequence | Region in protein | Ion Score |
|---|---|---|---|---|---|---|---|
| H6ST1 no TPST1/2 (DDA) | HS6ST1 | Heparan-sulfate 6-O-sulfotransferase 1 | 52 | 1530 | | | |
| | HS6T3 | Heparan-sulfate 6-O-sulfotransferase 3 | 22 | 548 | | | |
| H6ST1 +TSPST1/2 (DDA) | HS6ST1 | Heparan-sulfate 6-O-sulfotransferase 1 | 59 | 1599 | EDADEPGRVPTED**Y**MSHIIEK | 390-410 | 44 |
| | | | | | EDADEPGRVPTED**Y**MSHIIEKW | 390-411 | 28 |
| | HS6T3 | Heparan-sulfate 6-O-sulfotransferase 3 | 9 | 381 | | | |
| H6ST1 + TPST1/2 (NL) | HS6ST1 | Heparan-sulfate 6-O-sulfotransferase 1 | 7 | 120 | EDADEPGRVPTED**Y**MSHIIEK | 390-410 | 59 |
| | | | | | EDADEPGRVPTED**Y**MSHIIEKW | 390-411 | 51 |
| | | | | | EDADEPGRVPTED**Y**MSHIIEKW | 390-411 | 27 |
| | TPST1 | Protein-tyrosine sulfotransferase 1 | 9 | 61 | LG**Y**DP**Y**ANPPNYGKPDPK | 324-341 | 52 |
| H6ST2 no TPST1/2 (DDA) | HS6ST2 | Heparan-sulfate 6-O-sulfotransferase 2 | 30 | 627 | | | |
| | HS6ST3 | Heparan-sulfate 6-O-sulfotransferase 3 | 19 | 197 | | | |
| H6ST2 + TPST1/2 (DDA) | HS6ST2 | Heparan-sulfate 6-O-sulfotransferase 2 | 43 | 1395 | EQNDNTSNGTND**Y**IGSVEK | 585-603 | 95 |
| | GEMIN5 | Gem-associated protein 5 | 1 | 32 | V**Y**EAVELLK | 991-999 | 32 |
| | TPST1 | Protein-tyrosine sulfotransferase 1 | 53 | 2821 | LG**Y**DPYANPPNYGKPDPK | 324-341 | 48 |
| H6ST2 + TPST1/2 (NL) | HS6ST2 | Heparan-sulfate 6-O-sulfotransferase 2 | 3 | 105 | EQNDNTSNGTND**Y**IGSVEK | 585-603 | 106 |
| | TPST1 | Protein-tyrosine sulfotransferase 1 | 5 | 83 | LG**Y**DP**Y**ANPPNYGKPDPK | 324-341 | 42 |
| | TUBA1A | Tubulin alpha-1A chain | 6 | 48 | EDMAALEKD**Y**EEVGVDSVEGEGEEEGEEY | 423-451 | 48 |
| H6ST3 + TPST1/2 (DDA) | HS6ST3 | Heparan-sulfate 6-O-sulfotransferase 3 | 63 | 2453 | SPTPDELPTC**Y**PGDDWSGVSLR | 275-296 | 42 |
| | | | | | EDGAAEGTVTED**Y**NSQVVR | 452-470 | 59 |
| | | | | | DHQWPKEDGAAEGTVTED**Y**NSQVVR | 446-470 | 71 |
| | TPST2 | Protein-tyrosine sulfotransferase 2 | 60 | 3709 | VLKGD**Y**K | 348-354 | 33 |
| | TUBA1B | Tubulin alpha-1B chain | 65 | 1713 | LSVD**Y**GKK | 157-164 | 30 |
| | | | | | EDMAALEKD**Y**EEVGVDSVEGEGEEEGEEY | 423-451 | 44 |
| | TUBA1C | Tubulin alpha-1C chain | 60 | 1249 | LSVD**Y**GKK | 157-164 | 30 |
| | HNRNPH1 | Heterogeneous nuclear ribonucleoprotein H | 22 | 225 | DLN**Y**CFSGMSDHR | 263-275 | 31 |
| H6ST3 + TPST1/2 (NL) | HS6ST3 | Heparan-sulfate 6-O-sulfotransferase 3 | 10 | 475 | DHQWPKEDGAAEGTVTED**Y**NSQVVR | 446-470 | 104 |
| | | | | | EDGAAEGTVTED**Y**NSQVVR | 452-470 | 77 |
| | | | | | SPTPDELPTC**Y**PGDDWSGVSLR | 275-296 | 43 |
| | IRS4 | Insulin receptor substrate 4 | 1 | 128 | EADSSSD**Y**VNMDFTK | 914-928 | 49 |
| | | | | | EADSSSD**Y**VNMDFTKR | 914-929 | 97 |
| | TUBB | Tubulin beta chain | 8 | 108 | ISV**YY**NEATGGK | 47-58 | 65 |
| | | | | | NSS**Y**FVEWIPNNVK | 337-350 | 59 |
| | TUBA1A | Tubulin alpha-1A chain | 8 | 69 | EDMAALEKD**Y**EEVGVDSVEGEGEEEGEEY | 423-451 | 50 |
| | | | | | LSVD**Y**GKK | 157-164 | 39 |
| | TPST1 | Protein-tyrosine sulfotransferase 1 | 5 | 55 | LG**Y**DP**Y**ANPPNYGKPDPK | 324-341 | 47 |
| | HNRNPH1 | Heterogeneous nuclear ribonucleoprotein H | 3 | 43 | DLN**Y**CFSGMSDHR | 263-275 | 43 |
